# Biosynthetic heme of malaria parasite induces cerebral pathogenesis by regulating hemozoin formation and griseofulvin can prevent cerebral malaria

**DOI:** 10.1101/2021.04.27.441715

**Authors:** Manjunatha Chandana, Aditya Anand, Sourav Ghosh, Subhashree Beura, Sarita Jena, Amol Ratnakar Suryawanshi, Govindarajan Padmanaban, Viswanathan Arun Nagaraj

## Abstract

Heme-biosynthetic pathway of malaria parasite is dispensable for asexual stages, but essential for sexual and liver stages. Despite having backup mechanisms to acquire hemoglobin-heme, pathway intermediates and/or enzymes from the host, asexual parasites express heme pathway enzymes and synthesize heme. Here we show heme synthesized in asexual stages promotes cerebral pathogenesis by enhancing hemozoin formation. Hemozoin is a parasite molecule associated with inflammation, aberrant host-immune responses, disease severity and cerebral pathogenesis. The heme pathway knockout parasites synthesize less hemozoin, and mice infected with knockout parasites are completely protected from cerebral malaria and death due to anaemia is delayed. Biosynthetic heme regulates food vacuole integrity and the food vacuoles from knockout parasites are compromised in pH, lipid unsaturation and proteins, essential for hemozoin formation. Targeting parasite heme synthesis by griseofulvin - a FDA-approved drug, prevents cerebral malaria in mice and provides a new adjunct therapeutic option for cerebral and severe malaria.

## INTRODUCTION

Malaria remains a major concern of morbidity and mortality, especially with the emerging parasite resistance to artemisinin-based combination therapies (ACTs) and mosquito resistance to insecticides. According to World Health Organization (WHO), 229 million cases and 409,000 malaria deaths occurred in 2019^1^. Of the five *Plasmodium* species causing human malaria, *Plasmodium falciparum* (*Pf*) is the deadliest one responsible for more than 90% of the infections. The clinical manifestations of *Pf* malaria vary from mild (uncomplicated malaria) to severe (complicated malaria). Uncomplicated malaria is characterized by fever, headache, nausea, chills and mild anaemia. Complicated malaria is categorized by the existence of at least one criterion of disease severity that includes respiratory distress, metabolic acidosis, pulmonary edema, severe anaemia, jaundice, renal failure or neurological complications like impaired consciousness, convulsions etc. The typical outcome of severe malaria is multi-organ failure and/or cerebral malaria (CM) of which, CM is the most severe neurological complication with high mortality. About one-third of the patients recovering from CM show long-term neurocognitive impairments^2–4^.

Our current understanding on CM comes from a few post-mortem studies of human CM (HCM) and a large number of experimental CM (ECM) studies performed in mouse models^5, 6^. There occurs increased permeability of blood-brain barrier (BBB), brain capillary occlusions, parasitized-red blood cells (pRBCs) accumulation in brain microvasculature, and endothelial activation with dysregulated inflammation and aberrant host-immune responses^7,8,9,10,11,12,13,14,15^ that lead to BBB disruption, intracerebral hemorrhages, ischemia, edema, increased intracranial pressure, axonal damage and demyelination, culminating in the dysfunction of central nervous system^16,17,18,19,20,21,22,23^. Cerebral pathology arises due to a complex interplay of molecular events triggered by various host- and parasite-derived factors. The synchronous growth of the asexual parasites in red blood cells (RBCs) and the associated schizogony result in the release of pathogen-associated molecular patterns (PAMPs) such as hemozoin (Hz), glycosylphosphatidylinositiol, parasite DNA and RNA, and danger-associated molecular patterns (DAMPs) such as heme, uric acid, and microvesicles^14, 15^. Hz and its precursor heme play a central role in CM pathogenesis^14, 24,25,26,27,28,29,30,31^. The asexual stage parasites endocytose host hemoglobin (Hb) and digest it in the food vacuole (FV). The toxic free heme released during this process is detoxified into Hz by heme detoxification protein (HDP) that undergoes a circuitous trafficking and abundantly present in infected RBCs than parasite FVs^32^. Although the rate of HDP-mediated Hz formation is much higher^32^, autocatalytic^33^, histidine rich protein (HRP)-mediated^34^ and lipid-driven mechanisms^35, 36^ for Hz formation have also been described. There is a positive correlation of Hz released into the circulation and phagocytosed by the circulating phagocytic cells with disease severity in children and adults^37,38,39^. Similarly, plasma free heme is associated with disease severity^40, 41^. Free heme is extremely cytotoxic to endothelial cells and it can increase the expression of adhesion molecules, induce NLRP3 inflammasome and IL-1β secretion, and activate polymorphonuclear cells. The only treatment option for CM is parenteral administration of artemisinin derivatives or quinine with supportive therapies. However, the fatality due to CM remains high despite the parasite clearance^2, 4, 42^. The molecular mechanisms underlying CM pathogenesis need to be understood for developing adjunct therapies.

Malaria parasite synthesizes heme *de novo* despite the ability of asexual stages to access host Hb-heme^43^. The parasite heme pathway is compartmentalized in mitochondrion, apicoplast and cytosol, and heme is eventually synthesized in the mitochondrion. Our earlier study with *P. berghei* (*Pb*) aminolevulinate synthetase (ALAS) and ferrochelatase (FC) knockouts (KOs) generated for the first and last enzymes, demonstrated that the parasite pathway is dispensable for asexual stages, but essential for the development of sporozoites in mosquitoes and pre-erythrocytic stages in liver. Moreover, FCKO parasites can utilize host Hb-heme for their survival in blood stages^44, 45^. Subsequent studies in *Pf* using the KO parasites generated for ALAS, FC, apicoplast-localized porphobilinogen deaminase and cytosol-localized coproporphyrinogen oxidase, confirmed these findings with the suggestion that extracellular ALA acquired through new permeability pathways may lead to parasite heme synthesis^46, 47^. The conversion of extracellular ALA into protoporphyrin IX occurs in RBCs with the help of host enzymes, and heme is synthesized by parasite FC. Although *de novo* heme synthesis is non-essential and the asexual KO parasites can acquire heme/heme precursors from host RBCs and import some of the host enzymes, the parasite enzymes are expressed and heme synthesis occurs as evident from ^13^C/^14^C-ALA metabolic labelling studies^43,44,45,46,47,48,49,50,51,52,53,54,55,56^. We hypothesized that the parasite *de novo* heme pathway should have a physiological relevance in the asexual stages. Here, we demonstrate that *de novo* heme induces CM pathogenesis by regulating Hz formation in the asexual stages, thus offering an answer to a long-standing question. We further provide a therapeutic option on the basis that griseofulvin - a well-known FDA-approved drug capable of inhibiting parasite heme synthesis, can prevent CM in mice.

## RESULTS

### Mice infected with heme pathway KO parasites are protected from CM

Our earlier work with *Pb* heme pathway KO parasites was carried out in outbred Swiss mice that do not develop CM^44^. Here, we performed our studies in CM-susceptible C57BL/6 inbred mouse strain by injecting 10^5^ asexual stage parasites. Assessment of the peripheral blood parasitemia showed 2-3 days delay in the growth of KO parasites with respect to the WT (Fig. 1a). Importantly, about 80% of the WT-infected mice succumbed to CM within day 10 when the blood parasitemia was around 10-30%. The WT-infected mice that escaped from CM died of anaemia on day 12-16 post-infection. In contrast, mice infected with ALASKO and FCKO parasites were completely protected from CM and they died because of anaemia on day 20-30 (Fig. 1b). The delay in the growth of KO parasites was associated with an early increase in the spleen weight of infected mice (Fig. 1c), suggesting a better splenic clearance. A similar delay in the growth of KO parasites and the mortality of KO-infected mice due to anaemia was observed in Balb/c mice that do not develop CM (Supplementary Fig. 1a,b). To rule out the possibility that CM protection is because of a delay in the increase in blood parasitemia, we performed growth analyses in C57BL/6 mice infected with 10^7^ ALASKO/FCKO parasites. While the growth of 10^7^ KO parasites in mice was comparable with 10^5^ WT parasites, KO-infected mice were once again completely protected from CM (Fig. 1d,e). The mortality due to anaemia was delayed by 6-10 days, and KO-infected mice could sustain a higher parasitemia for a prolonged period. There were no significant differences in the reticulocyte versus mature RBC preference between WT and KO parasites in the first 9 days, the duration in which CM mortality occurred in WT-infected mice. However, KO parasites showed significantly increased reticulocyte preference and multiple infections in the reticulocytes during the later course of infections that represent anaemic phase (Fig. 1f,g). The rapid murine coma and behavioural scale (RMCBS) score of 10^5^ WT parasite-infected mice that succumbed to CM was below 5 on day 7 whereas, RMCBS score of 10^7^ KO parasite-infected mice was around 17 and 14 on day 7 and 14, respectively (Fig. 1h). For the subsequent experiments, we initiated asexual infections by injecting 10^5^ WT and 10^7^ ALASKO/FCKO parasites. These results were confirmed with another set of independent KO parasite lines wherein, ALAS and FC genes were replaced individually with GFP-luciferase (Luc)-expressing cassette containing m-cherry (Fig. 2a). The successful replacement of ALAS and FC was confirmed by PCR analyses performed with DNA and RNA isolated from the respective KO parasites (Fig. 2b,c), and by examining GFP and m-cherry fluorescence (Fig. 2d). For control, *c/d ssurRNA* locus in the WT parasite was replaced with GFP-Luc-expressing cassette containing m-cherry. *In vivo* bioluminescence studies showed accumulation of pRBCs in the brain of WT-infected mice, but not in the KO-infected mice (Fig. 2e) and this was confirmed by *ex vivo* imaging as well (Fig. 2f). The mice infected with Luc-expressing KO parasites were also completely protected from CM (Fig. 2g). These data indicated that the mice infected with heme pathway KO parasites are completely protected from CM.

**Fig. 1.**
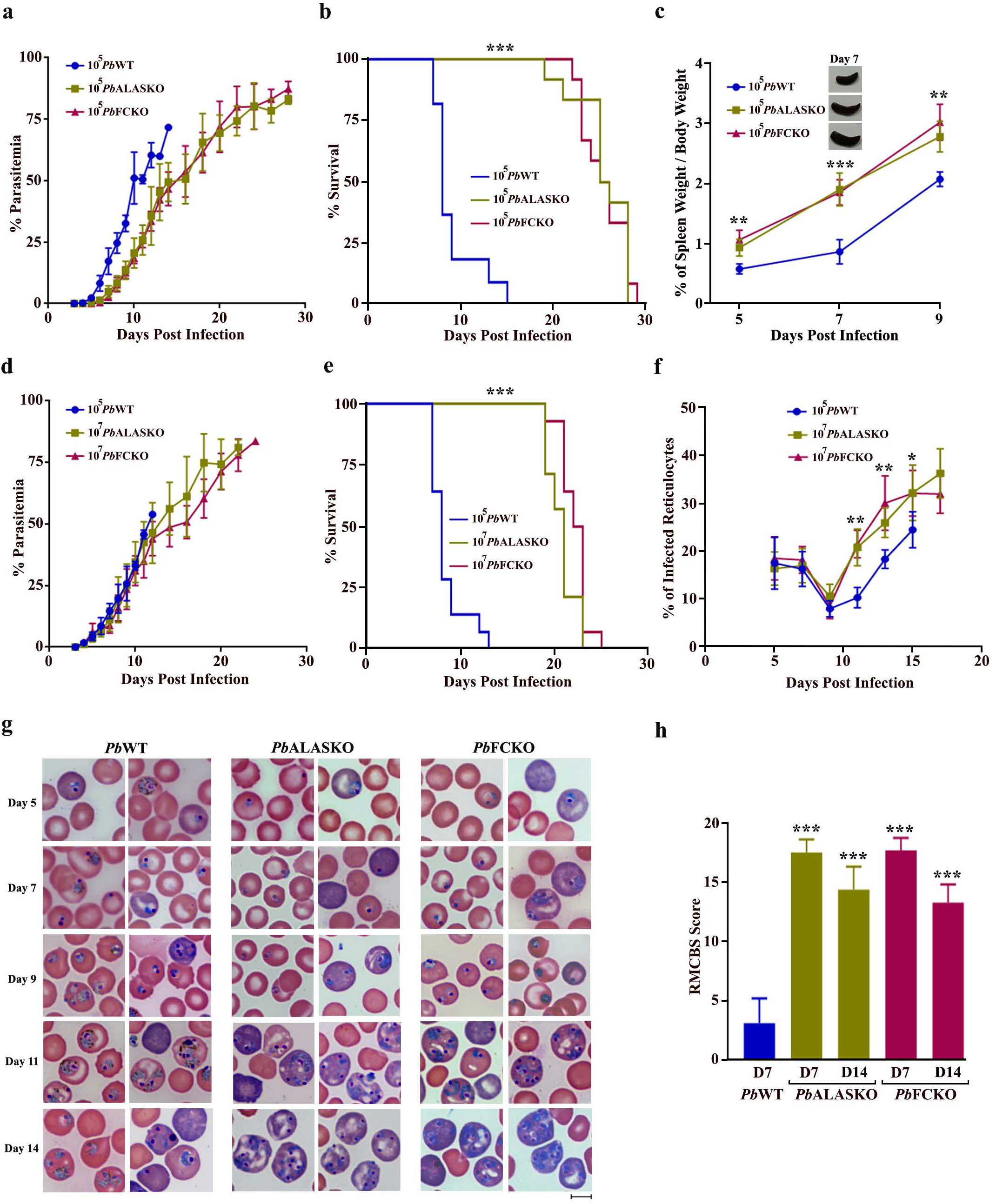
CM protection in heme pathway KO parasite-infected mice. **a**, Growth analysis of *Pb*WT (n=10), *Pb*ALASKO (n=12) and *Pb*FCKO (n=12) parasites in C57BL/6 mice. 10^5^ parasites were used to initiate *Pb*WT and *Pb*KO parasite infections. The data represent three different batches. **b**, Mortality curves of mice infected with *Pb*WT, *Pb*ALASKO and *Pb*FCKO parasites. The data represent the mice utilized for growth curve analysis (\*\*\**P*<0.001, log-rank (Mantel-Cox) test). **c**, Spleen weight of mice infected with *Pb*WT (n=13), *Pb*ALASKO (n=14) and *Pb*FCKO (n=14) parasites. For each day, 3-5 mice from four different batches were included (mean ± SD \*\**P*<0.01, \*\*\**P*<0.001, unpaired t-test). **d**, Growth analysis of *Pb*WT (n=14), *Pb*ALASKO (n=14) and *Pb*FCKO (n=14) parasites in C57BL/6 mice. 10^5^ and 10^7^ parasites were used to initiate WT and KO parasite infections, respectively. The data represent four different batches. **e**, Mortality curves of mice infected with *Pb*WT, *Pb*ALASKO and *Pb*FCKO parasites. The data represent the mice utilized for growth curve analysis (\*\*\**P*<0.001, log-rank (Mantel-Cox) test). **f**, Percentage of infected reticulocytes in the parasitized red cells (\**P*<0.05, \*\**P*<0.01, unpaired t-test). The data represent six mice each for *Pb*WT, *Pb*ALASKO and *Pb*FCKO parasites. **g**, Giemsa-stained images for peripheral blood smears prepared from tail vein blood of *Pb*WT and *Pb*KO parasite-infected mice. Reticulocytes could be identified by their distinct blue color. Images were captured using 100x objective lens. Scale bar = 5 μm. **h**, RMCBS score for mice infected with *Pb*WT (n=8) and *Pb*KO (n=12) parasites (mean ± SD; \*\*\**P*<0.001, unpaired t-test).

**Fig. 2.**
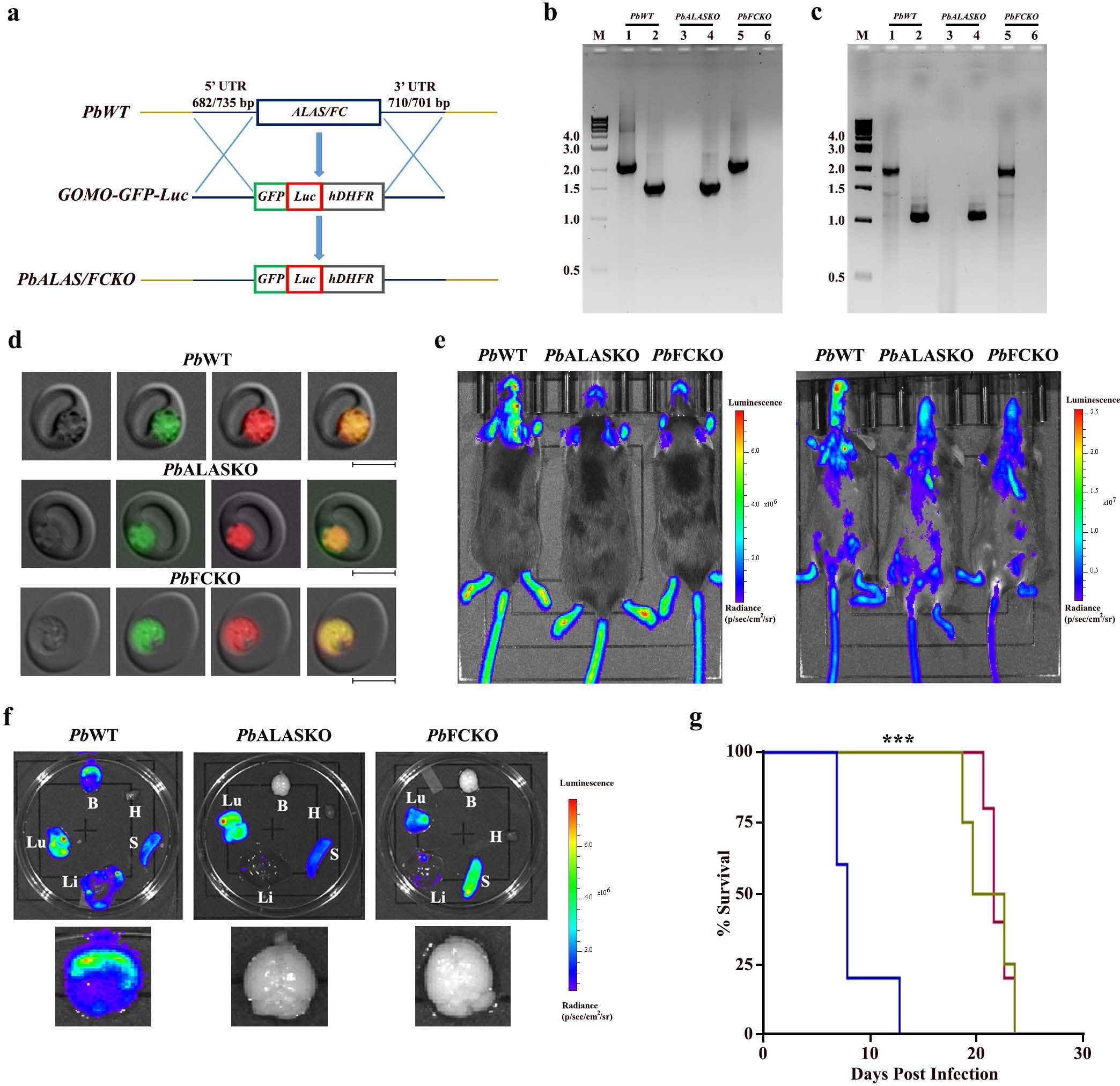
Generation of Luc-expressing heme pathway KO parasites and *in vivo* bioluminescence imaging of infected mice. **a**, Double crossover recombination strategy utilized to generate Luc-expressing *Pb*ALASKO and *Pb*FCKO parasites. **b**, PCR confirmation for *ALAS* and *FC* deletions with genomic DNA isolated from Luc-expressing *Pb*ALASKO and *Pb*FCKO parasites. Lane M: 1 kb ladder; Lane 1, 3 and 5: ALAS product (2.11 kb); Lane 2, 4 and 6: FC product (1.54 kb). **c**, RT-PCR confirmation for *ALAS* and *FC* deletions with total RNA isolated from Luc-expressing *Pb*ALASKO and *Pb*FCKO parasites. Lane M: 1 kb ladder; Lane 1, 3 and 5: ALAS product (1.92 kb); Lane 2, 4 and 6: FC product (1.05 kb). **d**, Live GFP and m-cherry fluorescence of Luc-expressing *Pb*WT and heme pathway *Pb*KO parasites. Images were captured using 100x objective lens. Scale bar = 5 μm. **e**, Whole body bioluminescence imaging of WT and KO parasite-infected mice on day 8 post-infection. **f**, *Ex vivo* bioluminescence imaging of liver (Li), lungs (Lu), brain (B), heart (H) and spleen (S) of Luc-expressing *Pb*WT- and *Pb*KO-infected mice. Enlarged images of brain are shown. **g**, Mortality curves of mice infected with Luc-expressing *Pb*WT (n=5), *Pb*ALASKO (n=4) and *Pb*FCKO (n=5) parasites (\*\*\**P*<0.001, log-rank (Mantel-Cox) test).

### Absence of cerebral pathology in heme pathway KO parasite-infected mice

To evaluate the integrity of BBB and assess vascular leakage, Evans blue extravasation analyses were carried out. While the brain collected from WT-infected mice on day 7/8 stained intensely with Evans blue, the extravasation of Evans blue into the brain was barely detectable in KO-infected mice on day 7 and 14 (Fig. 3a). Quantification of Evans blue in the brain extracts confirmed this observation (Fig. 3b). Histopathological assessment of hematoxylin and eosin (H&E)-stained brain sections of WT-infected mice on day 7 showed intracerebral hemorrhages with extravasation of erythrocytes into the perivascular space, petechial hemorrhages, thrombosed and leukocyte-packed vessels, gross demyelination and myelin pallor. No such hallmark features of CM could be detected in the brain sections of KO-infected mice (Fig. 3c). Immunohistochemical studies performed with the brain sections of WT-infected mice showed the extravasation of IgG in cerebral parenchyma and the presence of IgG in occluded vessels and hemorrhages, but not in the KO-infected mice (Fig. 3d). Luminal and abluminal leukocytes, and parasite-derived Hz could be detected in the occluded vasculature of WT-infected mice. (Fig. 3e). Immunofluorescence analyses of the brain sections using *Pb* glyceraldehyde-3-phosphate dehydrogenase (GAPDH) and mouse CD31 antibodies showed the accumulation of parasites in CD31^+^ vasculature and extravascular parasites in the hemorrhages of WT-infected mice, but not in the KO-infected mice (Fig. 3f). Similarly, antibodies specific for CD3 and β-amyloid precursor protein (β-APP) showed the accumulation of CD3^+^ T cells in the cerebral vasculature (Fig. 3g) and axonal injury in the brain sections of WT-infected mice, but not in the KO-infected mice (Fig. 3h). These data suggested the absence of CM-associated brain lesions in mice infected with heme pathway KO parasites.

**Fig. 3.**
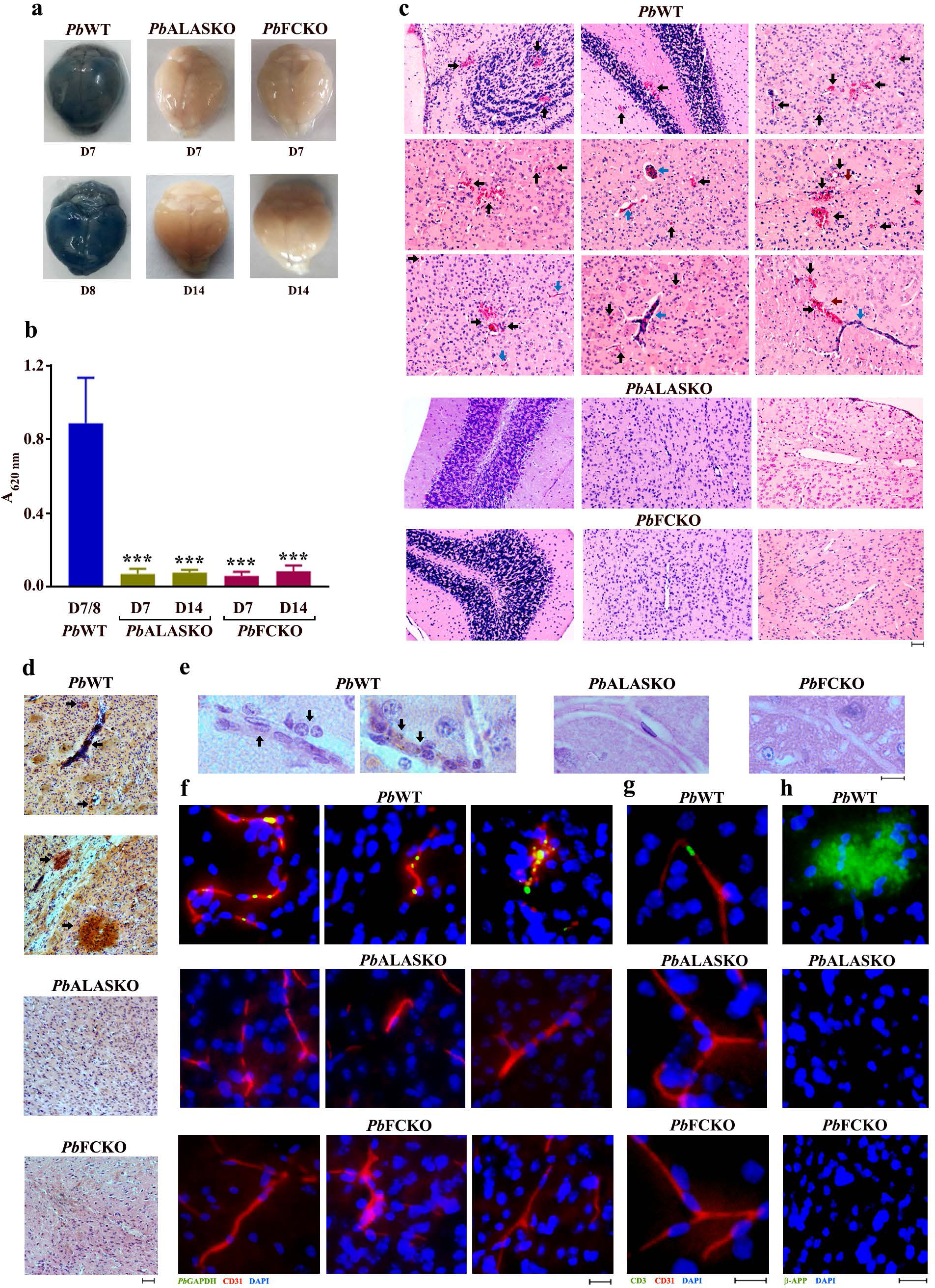
Assessment of cerebral pathology in heme pathway KO parasite-infected mice. **a**, Evans blue extravasation in the brain of mice infected with *Pb*WT and heme pathway *Pb*KO parasites. **b**, Quantification of Evans blue in the brain samples of mice infected with *Pb*WT (n=6) and heme pathway *Pb*KO (n=9) parasites. (mean ± SD) (\*\*\**P*<0.001, unpaired t-test). **c**, H&E staining of the brain sections prepared from *Pb*WT and *Pb*KO parasite-infected mice. Black arrows - intracerebral and petechial hemorrhages, blue arrows - thrombosed blood vessels and brown arrows - gross demyelination. Images were captured using 10x objective lens. Scale bar = 50 μm. **d**, IgG extravasation in the brain sections of *Pb*WT and *Pb*KO parasite-infected mice. Black arrows - areas showing IgG immunoreactivity. Images were captured using 10x objective lens. Scale bar = 50 μm. **e**, H&E staining of the brain sections indicating (black arrows) occluded vasculatures containing luminal and abluminal leukocytes, and parasite-derived Hz. Images were captured using 60x objective lens. Scale bar = 10 μm. **f**, Immunofluorescence analysis of parasite accumulation in the brain sections of *Pb*WT and *Pb*KO parasite-infected mice. **g**, Immunofluorescence analysis of CD3^+^ cells in the blood vessels of *Pb*WT and *Pb*KO parasite-infected mice. **h**, Immunofluorescence analysis of β- APP staining in the brain sections of *Pb*WT and *Pb*KO parasite-infected mice. Images were captured using 20x objective lens. Scale bar = 20 μm.

### Levels of inflammatory parameters in heme pathway KO parasite-infected mice

Multiplex assays performed for mice infected with KO parasites on day 8 showed a significant decrease in the plasma levels of proinflammatory cytokines and chemokines - IL- 6, TNFα, IFNγ, G-CSF, CCL3 (MIP-1α) and CCL5 (RANTES). There was also a significant increase in anti-inflammatory cytokines - IL-4, IL-10 and IL-13 (Fig. 4a). Quantitative RT-PCR analyses examining the expression of cytokines, chemokines, chemokine receptors and other key mediators of cerebral pathogenesis in the brain samples of KO-infected mice showed a substantial reduction of more than 1.5 fold in the transcript levels of TNFα, IFNγ, CXCL9, CXCL10, CCL2 (MCP-1), CCL5, CCL19, perforin, granzyme B, ICAM-1, p-selectin and HO-1. In particular, the decrease in IFNγ, CXCL10, CCL2, CCL5, granzyme B, ICAM-1 and HO-1 in FCKO was greater than 4-fold (Fig. 4b). Flow cytometry analyses of the leukocytes isolated from the brain samples of KO-infected mice showed significant reduction in CD3^+^ CD4^+^ and CD3^+^ CD8^+^ double positive cells, and CD3^+^ CD8^+^ CD69^+^, CD3^+^ CD8^+^ CXCR3^+^, CD3^+^ CD8^+^ perforin^+^, CD3^+^ CD8^+^ granzyme B^+^, CD3^+^ CD8^+^ TNFα^+^ and CD3^+^ CD8^+^ IFNγ^+^ triple positive cells (Fig. 4c and Supplementary Fig. 2). Western analyses for the brain homogenates of KO-infected mice showed reduction in phospho-NLRP3, phospho-NF-κB, cleaved caspase-1 and IL-1β (Fig. 4d). These data indicated an overall decrease in systemic and neuronal inflammation, and T cell infiltration in the brain milieu, suggesting that the cerebral pathology is impeded in mice infected with heme pathway KO parasites.

**Fig. 4.**
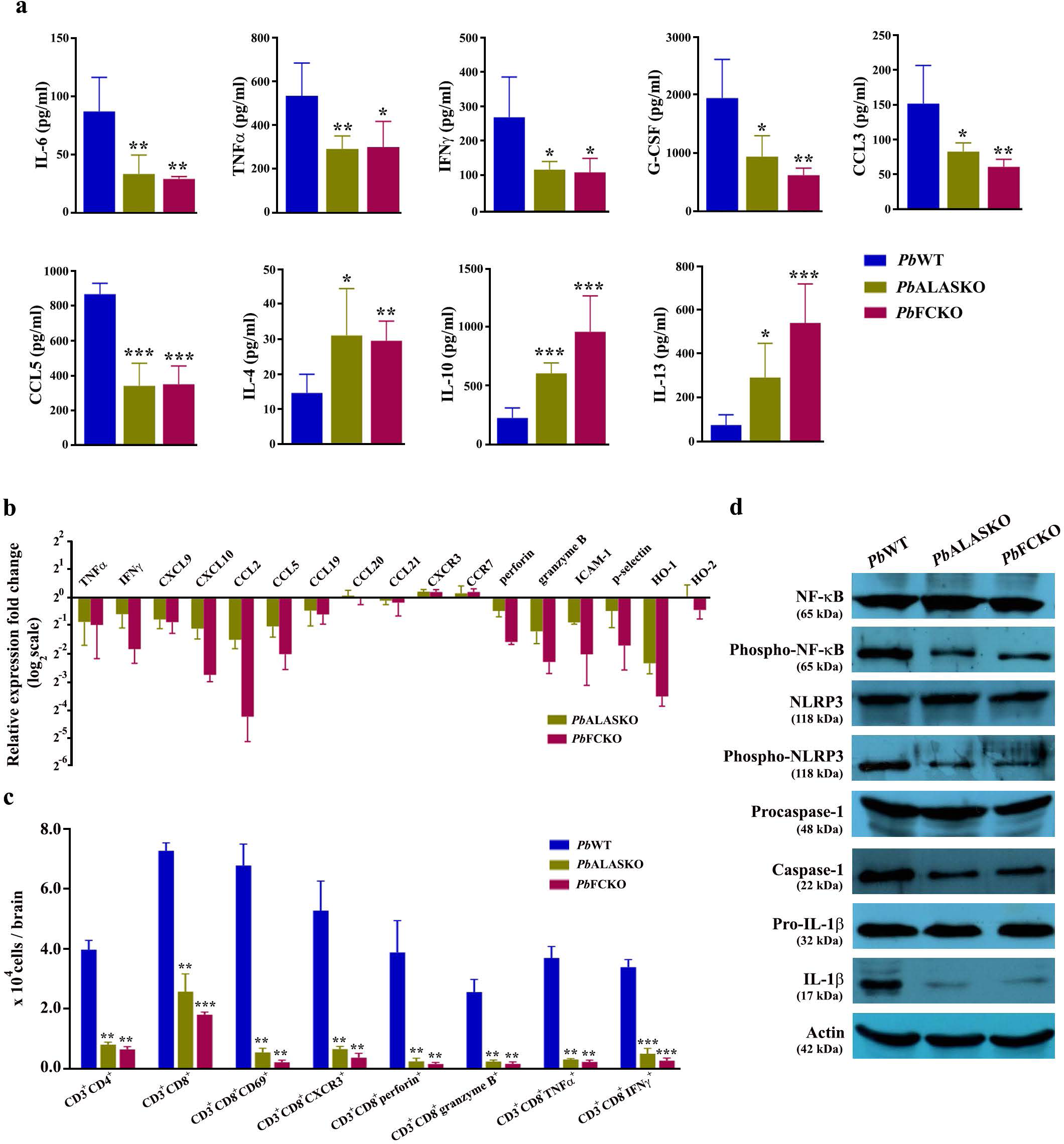
Assessment of inflammatory parameters in heme pathway KO parasite-infected mice. **a**, Plasma cytokine and chemokine levels of *Pb*WT- and *Pb*KO-infected mice (n=5) (mean ± SD; \**P*<0.05, \*\**P*<0.01, \*\*\**P*<0.001, unpaired t-test). **b**, qPCR analyses of host transcripts in the brain samples of infected mice. Expression levels were normalized with mouse GAPDH. Relative expression fold changes of mRNA transcripts in the KO-infected mice with respect to WT-infected mice (mean ± SD) are shown (n=3). **c**, Flow cytometry analyses of T cells in the brain samples of infected mice. Mice on day 7/8 post-infection were used and the data for each cell type were obtained from three different mice infected with *Pb*WT or *Pb*KO parasites (mean ± SD; \*\**P*<0.01, \*\*\**P*<0.001, unpaired t-test). **d**, Western analyses of brain homogenates prepared from *Pb*WT- and *Pb*KO-infected mice. 200 μg of total protein was used from the pooled brain homogenates of three different mice for *Pb*WT, *Pb*ALASKO and *Pb*FCKO.

### Decreased Hz formation in heme pathway KO parasites

A substantial decrease for Hz formation in the asexual stages of KO parasites was observed in the Giemsa-stained peripheral blood smears (Fig. 5a). This was more prominent in the asexual stages than gametocytes (Fig. 5b) and could be readily detected in almost 60-70% of the pRBCs containing asexual stages. These findings were verified by examining Hz content in paraformaldehyde-fixed pRBCs (Fig. 5c) and Hz dynamics in live pRBCs (Supplementary Movie 1, Supplementary Movie 2 and Supplementary Movie 3). To confirm, total Hz content of the parasites normalized with respect to the protein was examined and it was only 20-25% of the WT (Fig. 5d). During Hz formation in the FV, heme can leach into the parasite cytosol. Free heme in the parasite lysates of the KO parasites prepared by hypotonic lysis showed around 55% decrease when compared with WT (Fig. 5e). In addition, there was a significant decrease of around 55-60% in the plasma free heme and heme/hemopexin ratio of the KO-infected mice (Fig. 5f,g). However, there were no significant differences between the plasma hemopexin levels of WT- and KO-infected mice (Fig. 5h), and there was only a marginal 15-20% decrease in the plasma Hb levels that was statistically significant in case of FCKO, but not in ALASKO (Fig. 5i). The quantification of Hz load in the spleen and liver showed 40-50% decrease in the KO-infected mice (Fig. 5j,k). A similar decrease in the Hz content and free heme was observed for the KO parasites isolated from Balb/c mice (Supplementary Fig. 3a,b). These results suggested an overall decrease in the Hz synthesis of KO parasites.

**Fig. 5.**
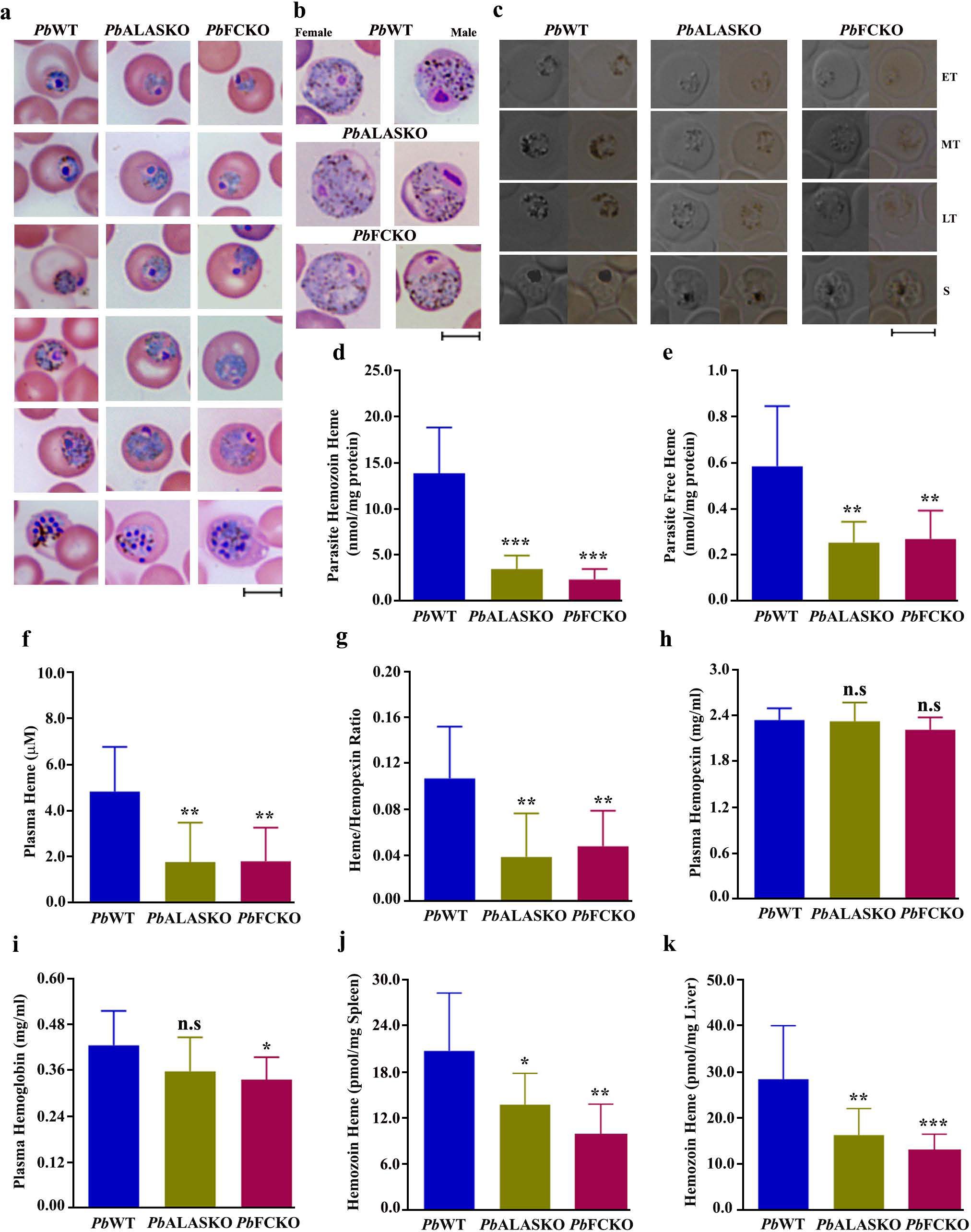
Hz and heme levels in KO parasites. **a,b**, Bright field images of Giemsa-stained PbWT, PbALASKO and PbFCKO asexual stage parasites and gametocytes, respectively, showing Hz content. Images were captured using 100x objective lens. Scale bar = 5 μm. **c**, Hz content in differential interference contrast (DIC; left) and bright field images (right) of paraformaldehyde-fixed pRBCs containing *Pb*WT and *Pb*KO parasites. Images were captured using 100x objective lens. Scale bar = 5 μm. **d**, Hz levels in *Pb*WT and *Pb*KO parasites **e**, Free heme levels in *Pb*WT and *Pb*KO parasites. **f**, Free heme levels in the plasma samples of *Pb*WT and *Pb*KO parasite-infected mice. **g**, Heme/Hemopexin ratio in the plasma samples of *Pb*WT and *Pb*KO parasite-infected mice. **h**, Plasma hemopexin levels of *Pb*WT and *Pb*KO parasite-infected mice. **i**, Plasma hemoglobin levels of *Pb*WT and *Pb*KO parasite-infected mice. **j,k**, Hz load in the spleen and liver of *Pb*WT and *Pb*KO parasite infected mice, respectively. **c-k**,. The data represent 9 mice each for *Pb*WT, *Pb*ALASKO and *Pb*FCKO. (mean ± SD; n.s - not significant, \**P*<0.05, \*\**P*<0.01, \*\*\**P*<0.001, unpaired t-test).

Heme and Hz are associated with the function of antimalarial drugs - artemisinin and chloroquine^57, 58^. Therefore, we were interested in examining the effect of α,β-arteether (artemisinin derivative) and chloroquine on FCKO-infected C57BL/6 mice *in vivo*. For α,β-arteether treatment, a single intramuscular dose of 1 mg/mouse or 0.4 mg/mouse was administered on day 5 post-infection when the blood parasitemia was around 5%. For chloroquine treatment, a daily dose of 25 mg/kg or 10 mg/kg was administered intraperitoneally for five consecutive days from day 5 post-infection. While 1 mg of α,β-arteether could completely clear the WT and FCKO parasites, 50% recrudescence was observed for 0.4 mg dosage in the WT- and FCKO-infected mice. The day of recrudescence was comparable between WT and FCKO parasites and the parasites were detectable in peripheral smears prepared on day 12 post-infection. However, the growth of FCKO parasites in mice showing recrudescence was delayed in the subsequent days and this was reflected in the mortality (Fig. 6a,b). The assessment of parasite load at the time of recrudescence on day 12 using *Pb*GAPDH primers showed that *C_t_* values normalized against mouse GAPDH for FCKO-infected mice were approximately 1.5 cycles more than WT-infected mice (Supplementary Fig. 4a-d), suggesting 3-fold less parasite load in FCKO-infected mice. In case of chloroquine, recrudescence was observed for WT and FCKO parasites with both the doses that were tested and importantly, FCKO parasites appeared 2-4 days earlier than WT (Fig. 6c,d). This was also confirmed by performing qPCR analysis on day 13 for 25 mg/kg and day 10 for 10 mg/kg doses. In both the cases, *C_t_* values obtained for the parasite load in FCKO-infected mice were approximately 4 cycles lower than WT-infected mice, suggesting 16-fold higher parasite load in FCKO-infected mice (Supplementary Fig. 4e-h). Further, one out of four and two out of four WT-infected mice died of CM in 25 mg/kg and 10 mg/kg treatment, respectively. Despite showing an early recrudescence, all the FCKO-infected mice died of anaemia and their mortality was delayed in comparison to WT-infected mice (Fig. 6d). These results indicated that chloroquine sensitivity is compromised in FCKO parasites while α,β-arteether sensitivity remains almost unaltered *in vivo* in mice.

**Fig. 6.**
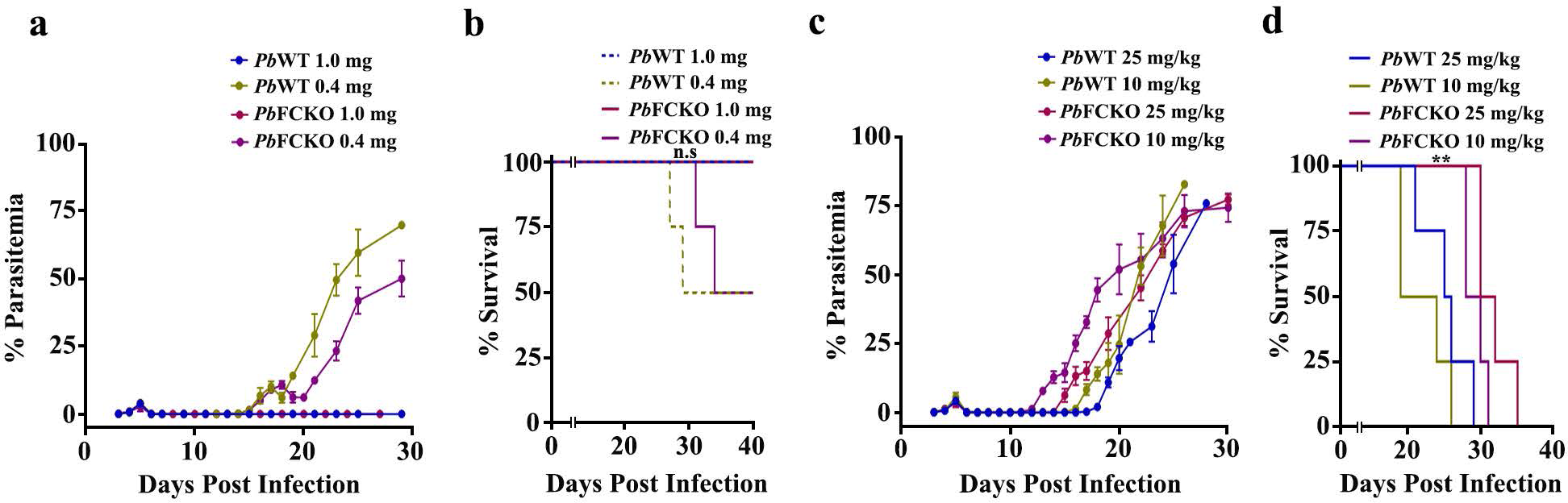
Sensitivity of WT and FCKO parasites to α,β-arteether and chloroquine. **a,b**, Blood parasitemia and mortality curves of infected mice treated with α,β-arteether, respectively. The data represent four mice for each group (n.s - not significant, log-rank (Mantel-Cox) test). **c,d**, Blood parasitemia and mortality curves of infected mice treated with chloroquine, respectively. The data represent four mice for each group (\*\**P*<0.01, log-rank (Mantel-Cox) test).

### *De novo* heme is essential for the functional integrity of FV

To understand the molecular mechanisms underlying decreased Hz synthesis, we assessed pH, lipids and protein content of the FVs in FCKO parasites and compared with WT parasites. Live fluorescence imaging of pRBCs incubated with acidophilic dye LysoTracker Deep Red indicated less fluorescence in the FCKO parasites (Fig. 7a), suggesting that the pH of FCKO FV is compromised. The quantification of fluorescence signal intensities showed ∼50% decrease (Fig. 7b). We examined the phospholipid content and could not find significant differences for *in vitro* ^32^P-orthophosphoric acid radiolabelling of major phospholipids - phosphatidylcholine (PC), phosphatidylinositol (PI) and phosphatidylethanolamine (PE), in the total parasites and FVs of FCKO parasites (Fig. 7c,d and Supplementary Fig. 5). Similarly, the assessment of neutral lipid content by live fluorescence imaging of pRBCs stained with BODIPY 493/503 and Nile Red did not show significant differences (Fig. 7e-h). Lipids associated with parasite Hz are reported to be abundant in monohydroxy derivatives of polyenoic fatty acids, and unsaturated glycerophospholipids induce rapid and efficient Hz formation in hematophagous insect *Rhodnius prolixus*^59, 60^. GC-MS analysis of fatty acid methyl esters (FAMEs) prepared from the FVs of *Pb* parasites indicated the presence of oleic acid (OA) as a major unsaturated fatty acid followed by arachidonic acid along with other saturated fatty acids, fatty acyl alcohols and derivatives, alkanes, etc. (Supplementary Fig. 6a and Supplementary Table 1). Malaria parasite scavenges stearic acid (SA) from plasma and converts it into OA by ER-localized Δ9-desaturase (stearoyl-CoA 9-desaturase). The *cis* double bond formation in OA requires heme since it utilizes the electrons transferred by cytochrome b5 from NADH cytochrome b5 reductase^61, 62^. *In vitro* ^14^C-SA radiolabelling and the separation of unsaturated FAMEs prepared from FCKO total parasites and FVs showed almost 80-90% decrease in the levels of OA methyl ester (OAME) in comparison to WT (Fig. 8a-c), with no significant differences in the signal intensities of SA methyl ester (SAME). The identity of the slower migrating polyunsaturated FAME (PU-FAME) that also showed a significant decrease is yet to be established and there are no clear-cut evidences for the presence of other desaturases in the parasite. There was a significant decrease in the radiolabelling of FV, but not in the total parasites (Supplementary Fig. 6b).

**Fig. 7.**
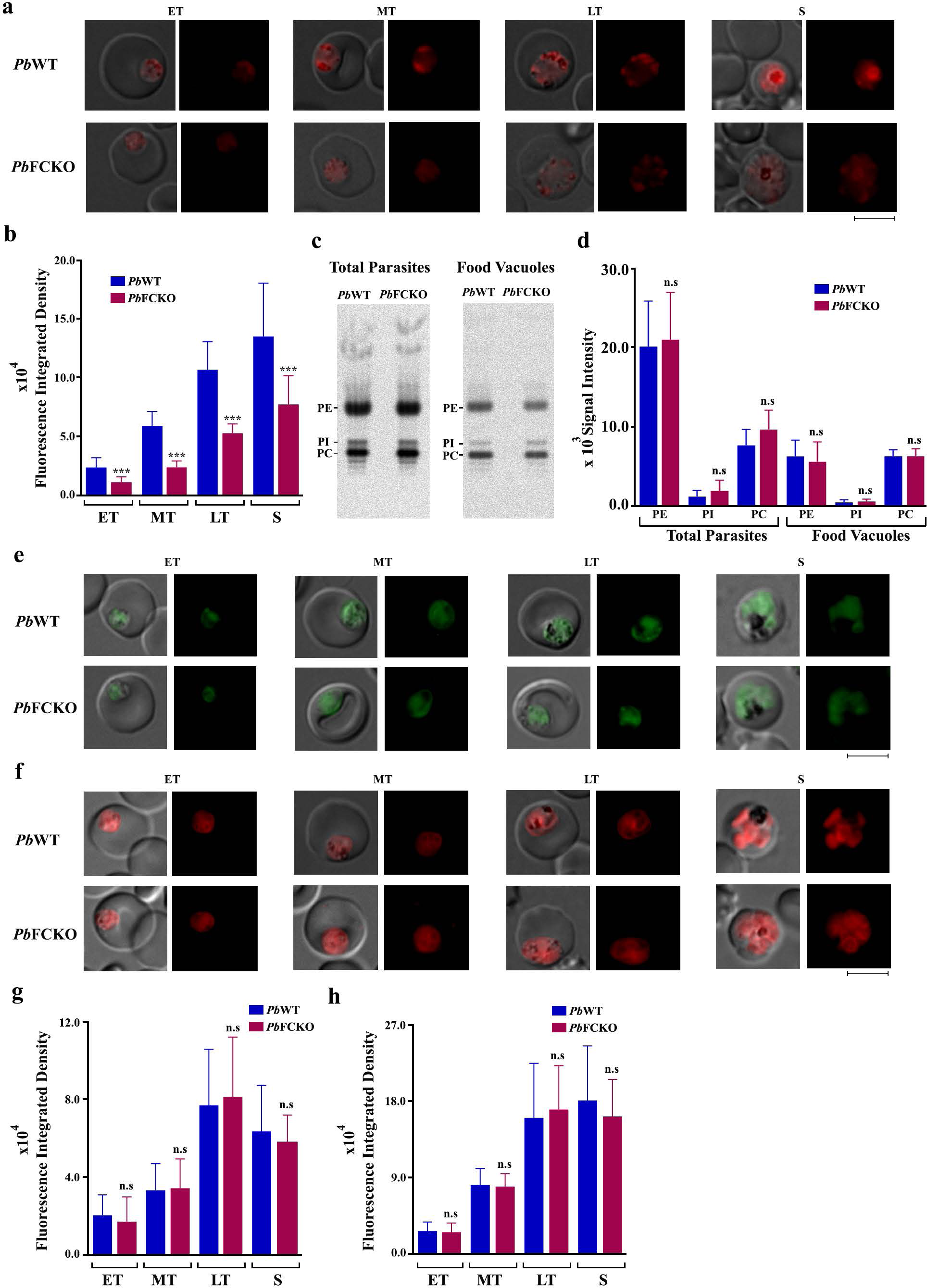
Assessment of food vacuole pH, phospholipids and neutral lipids in FCKO parasites. **a**, Live cell fluorescence imaging of LysoTracker Deep Red uptake in *Pb*WT and *Pb*FCKO parasites. **b**, Quantification of fluorescence signal from various stages. *Pb*WT - 98 trophozoites and 22 schizonts; *Pb*FCKO - 95 trophozoites and 23 schizonts. ET, MT and LT - early, mid and late trophozoites; S - schizonts. Images were captured using 100x objective lens. Scale bar = 5 μm. (mean ± SD; \*\*\**P*<0.001, unpaired t-test). **c**, TLC separation of ^32^P-orthophosphoric acid radiolabelled phospholipids for *Pb*WT and *Pb*FCKO total parasites and FVs. Half the phospholipid preparation was used for total parasites. For FVs, entire preparation was used. PC - phosphatidylcholine; PI - phosphatidylinositol; PE - phosphatidylethanolamine. **d**, Band intensities quantified (mean ± SD; n.s - not significant, unpaired t-test). The data represent three different experiments. **e**, Live cell fluorescence imaging of BODIPY 493/503 staining in WT and FCKO parasites. Images were captured using 100x objective lens. Scale bar = 5 μm. **f**, Live cell fluorescence imaging of Nile Red staining in WT and FCKO parasites. Images were captured using 100x objective lens. Scale bar = 5 μm. **g**, Quantification of the fluorescence signal from various stages for BODIPY 493/503 staining. The data represent 69 trophozoites and 15 schizonts for *Pb*WT, and 68 trophozoites and 14 schizonts for *Pb*FCKO (mean ± SD; n.s - not significant, unpaired t-test). **h**, Quantification of the fluorescence signal from various stages for Nile Red staining. The data represent 68 trophozoites and 14 schizonts for *Pb*WT, and 70 trophozoites and 14 schizonts for *Pb*FCKO (mean ± SD; n.s - not significant, unpaired t-test). ET - early trophozoites; MT - mid trophozoites; LT - late trophozoites; S - schizonts.

**Fig. 8.**
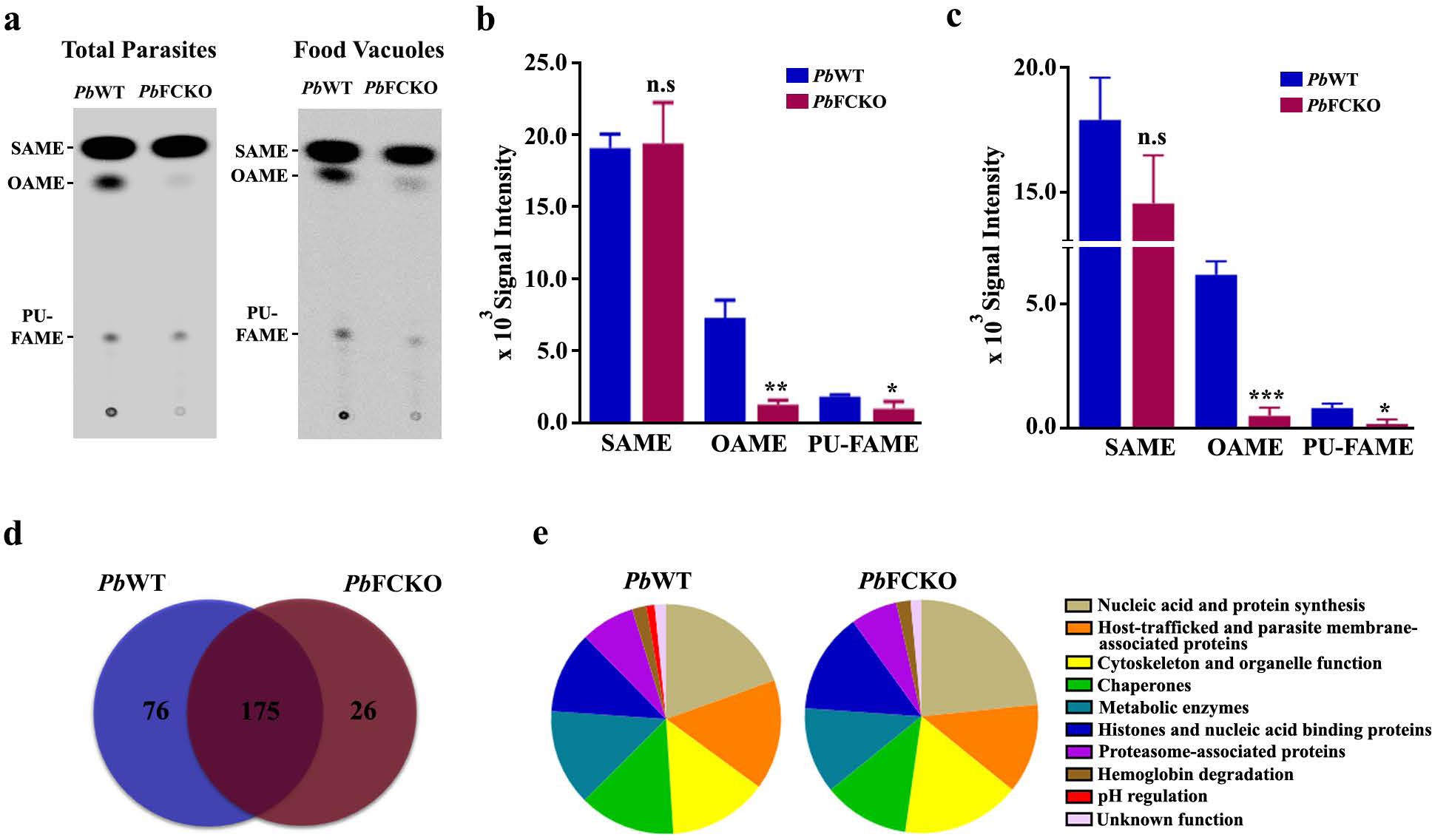
Evaluation of OA synthesis and FV proteomics for WT and FCKO parasites. **a**, TLC separation of ^14^C-SA radiolabelled unsaturated FAMEs for *Pb*WT and *Pb*FCKO total parasites and FVs. One third of the FAME preparation used for total parasites and entire preparation used for FVs. **b,c**, Band intensities quantified for total parasites and FVs, respectively (mean ± SD; n.s - not significant, \**P*<0.05, \*\**P*<0.01, \*\*\**P*<0.001, unpaired t-test), from three different experiments. **d**, Venn diagram of total proteins identified in the *Pb*WT and *Pb*FCKO FVs. **e**, Functional classification of the proteins based on gene ontologies available at PlasmoDB and UniProt databases. Supplementary Table 2 and Supplementary Table 3 have the complete set of details related to *Pb*WT and *Pb*FCKO FV proteome analyses.

FV proteomics is challenging due to the presence of large amounts of host Hb, Hb degradation peptides and indigenous proteases. There is only one report on *Pf* FV proteomics based on LC-MS/MS analysis of proteins excised from SDS-PAGE gel that identified 116 proteins excluding elongation factors and ribosomal proteins^63^. Our repeated attempts to perform differential proteomics for WT and FCKO FVs by Isobaric tags for relative and absolute quantitation (iTRAQ) were unsuccessful because of the interference from host Hb. LC-MS/MS of in-solution trypsin digested FV protein extracts solubilized with 6M urea was successful. The quality of FV preparations could be assessed by the presence of signature FV proteins such as plasmepsin IV, berghepain, aminopeptidases, subunits of vacuolar-type H^+^ATPase (V-type H^+^ ATPase), together with parasitophorous vacuolar (PV) proteins including exported protein 1 (Exp1), Exp2, early transcribed membrane protein, PV1, PV5 (lipocalin) etc., and Rab GTPases associated with cytostome-FV trafficking. A total number of 251 and 201 proteins could be identified for WT and FCKO FVs, respectively, and 175 proteins were common between them suggesting an overall consistency in the preparations (Fig. 8d). Around 68% of the proteins identified in the total *Pf* FV proteome were present in our preparations. Remarkably, in agreement with the decreased uptake of LysoTracker Deep Red in FCKO pRBCs, none of the subunits of V-type H^+^ATPase - a proton pump maintaining the acidic pH of FV^64^ could be detected in FCKO FVs indicating the less abundancy of these proteins. In WT FV preparations, A, B and G subunits of V-type H^+^ATPase could be detected. While plasmepsin IV, the only Pb aspartic protease involved in Hb degradation could be detected in WT and FCKO FVs, berghepain-2 - a cysteine protease involved in Hb degradation^65, 66^ could not be detected in FCKO FVs (Fig. 8e, Supplementary Table 2 and Supplementary Table 3). The rest of the proteins unique for WT and FCKO FVs include proteasome subunits, ribosomal proteins, metabolic enzymes etc., (Supplementary Table 2 and Supplementary Table 3) that are not related to Hz formation and known to be present in FV preparations^63^. All these evidences suggested that the functional integrity of FCKO FVs is compromised in terms of pH, lipid unsaturation and proteins that in turn can collectively lead to a decreased Hz formation.

### Griseofulvin treatment protects mice from ECM

Griseofulvin, isolated from *Penicillium griseofulvum*, is a FDA-approved antifungal drug used to cure tinea infections. It interacts with fungal microtubules and disrupts spindle assembly leading to mitotic arrest. In humans, griseofulvin dosage is given to the extent of 1000 mg/day in adults and 10 mg/kg/day in children for several weeks^67,68,69^. It can also inhibit FC by generating N-methyl protoporphyrin IX (NMPP) through the action of cytochrome P450 enzymes^70^. Therefore, we evaluated the potential of griseofulvin in preventing CM by treating WT-infected C57BL/6 mice from day 4 when the blood parasitemia was around 2%. A single dose of 2 mg/day (comparable with the dosage of humans) administered from day 4 and continued until day 8 showed the best protection. While ∼80% of the control mice succumbed to CM within day 10, more than 80% of the treated mice were protected from CM. Similar protection was also observed for the mice treated with 1 mg dose, twice a day from day 4 to day 8 (Fig. 9a). There was also a significant delay in the mortality that occurred due to anaemia. While the mortality in CM- escaped control mice occurred within day 17, more than 80% of the treated mice could survive beyond day 20 with almost 50% of them surviving even beyond day 24. Interestingly, the growth curves of WT parasites in treated and untreated mice were very much comparable (Fig. 9b). We analyzed heme synthesis in griseofulvin-treated WT parasites by incubating the *in vivo*-treated pRBCs *in vitro* with ^14^C-ALA for 9 h and there was around 60% decrease in the ^14^C-labelling of free heme (Fig. 9c). Further, griseofulvin-treated mice showed less Evans blue extravasation in the brain (Fig. 9d), with the absence of intracerebral hemorrhages (Fig. 9e), lack of accumulation of parasites and CD3^+^ T cells in the cerebral vasculature (Fig. 9f,g) and undetectable axonal injury (Fig. 9h). Giemsa-stained peripheral smears and paraformaldehyde-fixed pRBCs showed less Hz content (Fig. 9i,j). As observed for KO parasites, the total Hz content in the griseofulvin-treated parasites was around 50-60% less when compared with untreated parasites (Fig. 9k) and there was close to 60% decrease in the free heme levels (Fig. 9l). Further, the plasma levels of heme, hemopexin and Hb together with heme/hemopexin ratio of the griseofulvin-treated WT-infected mice were comparable with the KO-infected mice (Fig. 9m-p). These results indicated the potential of griseofulvin in preventing CM through the inhibition of parasite heme synthesis.

**Fig. 9.**
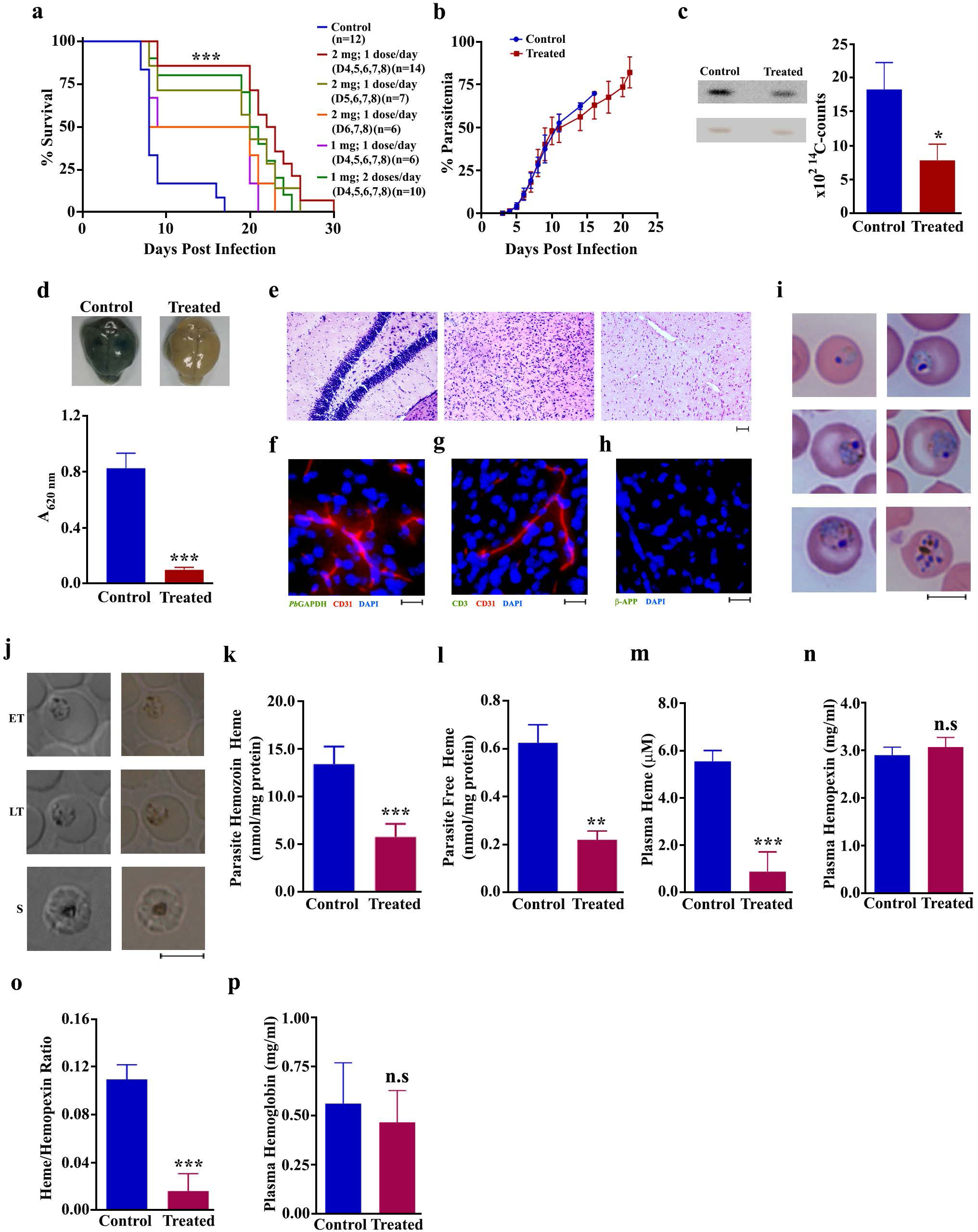
Effect of griseofulvin treatment on CM pathogenesis. **a**, Mortality curves of *Pb*WT- infected mice treated with different dosages of griseofulvin (\*\*\**P*<0.001, log-rank (Mantel-Cox) test). **b**, Growth curve analysis (n=12) for 2 mg dose per day on day 4,5,6,7 and 8. **c**, Phosphorimager and scanned images of TLC performed for ^14^C-ALA labelled parasite free heme and the radioactive counts measured for three different experiments (mean ± SD; n.s -not significant, \**P*<0.05, unpaired t-test). **d**, Extravasation of Evans blue and its quantification (n=3) (mean ± SD; \*\*\**P*<0.001, unpaired t-test). **e**, H&E staining of the brain sections. Images were captured using 10x objective lens. Scale bar = 50 μm. **f**, Parasite accumulation **g**, CD3^+^ cells in the blood vessels. **h**, Axonal injury in the brain sections. Images were captured using 20x objective lens. Scale bar = 20 μm. **i**, Giemsa-stained parasites. Images were captured using 100x objective lens. Scale bar = 5 μm. **j**, Hz content in differential interference contrast (DIC; left) and bright field images (right) of paraformaldehyde-fixed RBCs. Images were captured using 100x objective lens. Scale bar = 5 μm. **k**, Parasite Hz (n=5). **l**, Parasite free heme (n=5). **m**, Plasma free heme (n=6). **n**, Plasma hemopexin (n=6). **o**, Plasma heme/hemopexin ratio. **p**, Plasma hemoglobin (n=11). **k-p**, (mean ± SD; n.s - not significant, \*\**P*<0.01, \*\*\**P*<0.001, unpaired t-test).

## DISCUSSION

Parasite heme pathway has remained enigmatic for more than two decades and the dichotomy between *de novo* heme and host-heme/porphyrin acquisition pathways in the asexual stages is obscure. Here, we provide an explanation as to why asexual parasite expresses “dispensable” heme pathway enzymes and synthesizes heme despite acquiring host heme, by demonstrating that *de novo* heme pathway is associated with disease virulence and it induces CM pathogenesis by promoting Hz formation. Hz is a key malarial PAMP associated with aberrant inflammatory responses, endothelial activation promoting pRBC sequestration, T cell infiltration and neuronal apoptosis. Upon phagocytosis, Hz can induce the production of pro-inflammatory cytokines and chemokines in neutrophils, monocytes, macrophages and DCs, leading to the increased expression of ICAM-1 in endothelial cells and enhanced sequestration of pRBCs. Hz can also trigger IL-1β production by activating NLRP3 inflammasome, and augment endothelial cell damage and loss of BBB integrity by inducing proinflammatory responses in endothelial cells and activating coagulation and complement pathways^14, 25^. The KO parasites synthesize less Hz and mice infected with KO parasites are completely devoid of cerebral complications. The plasma levels of IL-6, TNFα, IFNγ, G-CSF, CCL3 and CCL5 are significantly decreased in KO-infected mice. These cytokines and chemokines are known to be highly elevated in the serum samples of cerebral and severe malaria patients^25, 71, 72^. There is also a concomitant increase in anti-inflammatory cytokines such as IL-4, IL-10 and IL-13 in the KO-infected mice indicating an over-all decrease in the systemic inflammation.

The transcript levels of TNFα, IFNγ, CXCL9, CXCL10, CCL2, CCL5 and CCL19 are low in the brain samples of KO-infected mice. CXCL9 and CXCL10 are highly induced in the brain of ECM mice and elevated levels of CXCL10 in plasma and cerebrospinal fluid predict fatal CM in humans^73^. CCL2, CCL5 and CCL19 associated with leukocyte migration are known to be induced by Hz in monocyte-derived DCs, macrophages and neutrophils^14, 25, 26, 74^. Similar decrease was also observed in the transcript levels of ICAM-1, p-selectin, perforin, and granzyme B, the key adhesion and cytotoxic effector molecules. Low transcript levels of HO-1 in KO-infected mice are due to less inflammation and decreased extracellular heme - the two potent HO-1 inducers^29, 30^. Further, there is a decrease in the infiltration of CD8^+^ T cells expressing early activation marker CD69, and proinflammatory and cytotoxic effector molecules such as TNFα^+^, IFNγ^+^, CXCR3^+^, perforin and granzyme B that leads to endothelial leakage, BBB disruption and neuronal damage in CM. There is also a reduction in phospho-NF-κB and NLRP3 inflammasome formation with decreased caspase-1 activation and less production of IL-1β - a key mediator of inflammation, infiltration of immune cells and neuronal apoptosis.

The decrease in Hz synthesis is reflected in the reduced Hz load in the organs, and decreased free heme levels in the parasites and plasma samples of KO-infected mice. It is suggested that oxidized heme derived from host Hb release during schizont rupture contributes to free plasma heme responsible for severe malaria and CM pathogenesis^29,30,31^. Interestingly, plasma levels of Hb and heme scavenging proteins like hemopexin in KO-infected mice are comparable with WT-infected mice indicating that the decreased plasma heme levels are due to changes in parasite heme/Hz levels. While less Hz formation can be directly associated with reduced labile/free heme levels in KO parasites, the decrease in plasma free heme needs further investigations. Despite thriving in a heme-rich milieu, parasite maintains cytosolic free heme concentrations of ∼1.6 μM, comparable with ∼0.4-0.6 μM concentrations seen in mammalian cells^75^. It is possible that the parasite could have evolved with protein-/transporter-based mechanisms to dispose excess free heme from the cytosol and this could be the reason for reduced plasma free heme levels in KO-infected mice. For example, HRP capable of binding heme is secreted by the parasite. Altogether, our findings suggest that *de novo* heme in the blood stages can influence the levels of Hz and free heme - a key PAMP and DAMP associated with malaria pathogenesis. Chloroquine acts predominantly by inhibiting host Hb-heme polymerization into Hz and it has been suggested that activation of artemisinin requires heme^57, 58^. In agreement with decreased Hz formation, the sensitivity of FCKO parasites to chloroquine is compromised. However, the *in vivo* sensitivity to α,β-arteether remains unaltered and our results indicate that albeit a ∼50% decrease, the host-derived labile heme present in the cytosol of FCKO parasites is adequate to activate α,β-arteether and the role of *de novo* heme is confined to cerebral pathogenesis.

The host Hb degradation in asexual stage parasites can release as much as ∼15 mM heme^75^ and therefore, utilization of host heme cannot account for almost 75% decrease in the Hz synthesis of KO parasites. Here, we provide evidence for the unexpected role of *de novo* heme in influencing the detoxification of Hb-heme into Hz by regulating the FV integrity in asexual stages (Fig. 10). FVs of FCKO parasites are compromised in terms of pH, lipid unsaturation and proteins associated with Hz formation. It is known that acidic pH of FV is critical for Hb digestion by proteases and subsequent Hz formation. The decrease in LysoTracker Deep Red uptake indicate that the FCKO FVs are less acidic. Our results also indicate the less abundancy of V-type H^+^ATPase subunits and berghepain-2 in FCKO FVs. V-type H^+^ATPase is a proton pump mainly responsible for maintaining the acidic pH of FV and targeting V-type H^+^ATPase activity with concanamycin A or bafilomycin A1 can lead to the alkalinisation of FV and inhibition of parasite growth^64^. *Pb* has two isoforms of berghepain - berghepain-1 and -2 of which, berghepain-1 is associated with hepatic merozoite invasion and erythrocyte tropism, and berghepain-2 seems to be involved in Hb digestion^65, 66^. Importantly, we show drastic reduction in OA synthesis of FCKO parasites with not much change in phospholipids or neutral lipids, suggesting an alteration in the degree of lipid unsaturation that can affect Hz formation. OA synthesis in malaria parasite is catalysed by a heme-dependent, ER-localized, Δ9-desaturase and it does not occur in uninfected RBCs^61, 62^. Our results suggest the specific role of *de novo* heme in OA synthesis that cannot be compensated by host Hb-heme. The levels of unsaturated fatty acids are also shown to stimulate the function of V-type H^+^ATPase in plants^76^, and the decrease in OA synthesis and lipid unsaturation can affect other cellular processes such as membrane homeostasis, protein/lipid trafficking, protein folding, cell signalling etc^77^. Although host Hb-heme acquisition starts from late ring stages, it is evident that FV integrity and maturation depend on *de novo* heme. Our results provide new functional insights on *de novo* heme that could be a miniscule in comparison with massive amounts of heme derived from host Hb. Intriguingly, we show it is the *de novo* heme that influences the detoxification of host heme into Hz and disease pathogenesis. Despite the ability of blood stage parasites to manage host Hb-heme for survival, lack of *de novo* heme synthesis renders them less virulent. It would be interesting to examine whether (i) *de novo* heme regulates other metabolic pathways or organellar functions and influences the trafficking mechanisms associated with host-heme or Hb uptake and (ii) compartmentalization of *de novo* and Hb-heme make them functionally different. Further, the accessibility of Hb-heme to various organelles and its incorporation in hemoproteins may vary. Altogether, our findings suggest the importance of “dispensable” *de novo* heme pathway in the asexual stages despite its non-essentiality for parasite growth.

**Fig. 10.**
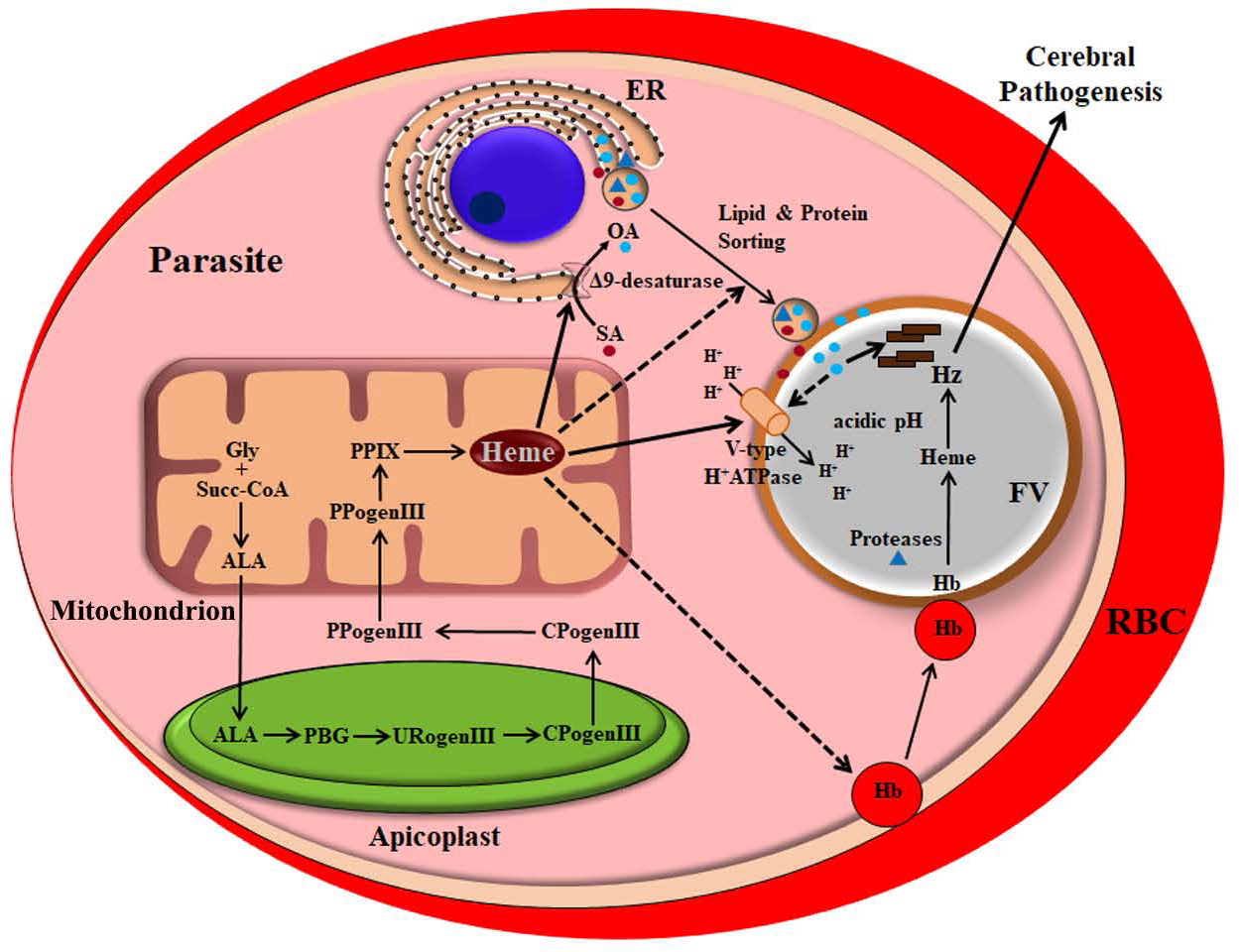
Model depicting the role of *de novo* heme in regulating FV integrity and Hz formation. *De novo* heme pathway of malaria parasite and Hz formation in the FV are represented. Solid arrows indicate the effect of *de novo* heme on V-type H^+^-ATPase, FV pH and OA synthesis affecting hemozoin formation. Dashed arrows represent the probable effects of *de novo* heme on Hb endocytosis, and protein and lipid trafficking. Gly - glycine; Succ-coA - succinyl coenzymeA; ALA - δ-aminolevulinic acid; PBG - prophobilinogen; URogenIII - uroporphyrinogen III; CPogenIII - coproporphyrinogen III; PPogenIII - protoporphyrinogen III; PPIX - protoporphyrin IX; ER - endoplasmic reticulum.

Finally, we show that *de novo* heme in the blood stages can serve as a target for malaria pathogenesis. Griseofulvin - a FDA-approved antifungal drug prevents CM in mice and delays death due to anaemia by inhibiting parasite heme synthesis. The ability of griseofulvin to prevent disease pathogenesis is observed despite the absence of any antimalarial treatment and griseofulvin treatment does not affect the parasite growth suggesting that CM protection is not because of the inhibition of parasite mitosis. This is in concurrence with a clinical trial conducted in malaria-infected humans where, griseofulvin treatment did not affect *in vivo* parasite growth although NMPP formation occurred within the parasite. The participants were rescued early with ACT and therefore, no observations were made on disease severity^78^. The fatality rates in malaria do not correlate with parasite clearance and therefore, targeting parasite virulence becomes important. Further, the malaria deaths that occur despite treating the infected individuals with ACTs and the decreasing efficacy of ACTs that lead to delayed parasite clearance underscore the need of an adjunct therapy in the initial stages of ACT treatment. Our study identifies a unique approach of targeting hemozoin through *de novo* heme to mitigate parasite virulence. Since *de novo* heme pathway is dispensable for asexual stages, there is a possibility of less selection pressure to result in resistance development. Repurposing of griseofulvin can serve as an excellent adjunct to ACTs for CM and severe malaria that needs to be evaluated in clinical trials.

## FUNDING

This study is supported by Centre of Excellence and Innovation in Biotechnology (CEIB) grant (BT/PR13760/COE/34/42/2015), Department of Biotechnology, New Delhi, (G.P. and V.A.N.), and intramural support from Institute of Life Sciences ILS/16-19 (V.A.N.), Bhubaneswar. G.P. is a NASI-Platinum Jubilee Senior Scientist.

## ACKNOWLEDGEMENTS

Thanks are due to Dr. Gulam Hussain Syed and Dr. Soumen Chakraborty for helping us with lipid labelling and transfection studies. We gratefully acknowledge the support rendered by Professor Balasubramanian Gopal, and mass spectrometry facilities at Molecular Biophysics Unit, Indian Institute of Science, Bangalore; Advanced Technology Platform Centre, Regional Centre for Biotechnology; and Kalinga Institute of Industrial Technology - Technology Business Incubator. We acknowledge the support rendered by Mr. R. Rajendra Reddy, Institute of Life Sciences Central Proteomics Facility, and Mr. Paritosh Nath, Institute of Life Sciences Flow Cytometry Facility. We thank Malaria Research and Reference Reagent Resource Center (MR4), ATCC Manassas Virginia, for providing us with *Pb* ANKA (MRA-311) deposited by Thomas F. McCutchan, and addgene for providing us with GOMO-GFP-LUC plasmid (#60976) deposited by Olivier Silvie.

## AUTHOR CONTRIBUTIONS

M.C., A.A., S.G., S.B. S.J. A.R.S. and V.A.N. performed the experiments. G.P. and V.A.N. conceived and designed the study. M.C., A.A., S.G., A.R.S., G.P. and V.A.N. analysed the data. M.C., A.A., S.G., G.P. and V.A.N. wrote the manuscript.

## COMPETING INTERESTS

The authors declare no competing financial interests.

## MATERIALS AND METHODS

### Routine propagation of *P. berghei* in mice and CM experiments

*P. berghei* ANKA WT and KO parasites were propagated in C57BL/6 male mice of 7-8 weeks old. Peripheral blood parasitemia was monitored by performing light microscopy for Giemsa-stained thin blood smears prepared from tail vein blood. When the blood parasitemia was around 10%, 10^5^ *P. berghei* ANKA WT or 10^5^/10^7^ ALAS/FC KO parasites were collected and injected intraperitoneally in 7-8 weeks old C57BL/6 male naïve mice to initiate CM experiments. CM phenotype was also confirmed with C57BL/6 female mice. For experiments carried out with Balb/c mice, 7-8 weeks old female mice were used. All the studies involving mice were carried out with the approval of Institutional Animal Ethics Committee (ILS/IAEC-57-AH/JAN-16) according to the national guidelines framed by “The Committee for the Purpose of Control and Supervision of Experiments on Animals (CPCSEA)”. Growth curve analysis was carried out by monitoring the blood parasitemia. The development and progression of ECM were monitored by examining RMCBS for neurological symptoms as described earlier^79^. To assess BBB integrity, Evans blue uptake assays were carried out by injecting 200 μl of 2% Evans blue in PBS intravenously and examining the extravasation of dye after one hour in the brain of the infected mice that were transcardially perfused with PBS. The extent of BBB damage was quantified by incubating the brain samples in formamide at 37°C for 48 h, extracting the Evans blue and measuring the absorbance at 620 nm^29^.

### Histological and immunofluorescence analyses of cerebral pathology in the brain of infected mice

For H&E staining to assess vascular blockage, hemorrhages and demyelination, brain samples were fixed with formalin for 72 hours at room temperature. After dehydrating with ethanol and treating with xylene, paraffin embedded blocks were made and sections of 7 μm thickness were prepared using Leica RM2125RT rotary microtome. The sections were then processed and stained with H&E using standard protocols. Immunohistochemical analysis of IgG extravasation in the brain sections was carried out as described^13^. In brief, brain sections of 30 μm thickness were antigen retrieved by treating them at 95°C for 30 min in sodium citrate buffer pH 6.0, followed by blocking with 3% H_2_O_2_ at room temperature for 30 min to prevent endogenous peroxidase activity. The sections were then incubated with HRP-conjugated goat anti-mouse IgG (Abcam, ab97023) at 1:250 dilution in PBS containing 0.3% Triton X-100 and 0.1% BSA, followed by developing with diaminobenzidine tertrahydrochloride (Vector Labs, SK-4100) and counterstaining with hematoxilin (HiMedia, S058). Immunoflourescence analysis of brain sections for parasite sequestration was carried out as described^13^ by fixing the brain samples in 4% paraformaldehyde in PBS containing 20% sucrose for 24 h at 4°C and cryoprotecting them for 48 h in PBS containing 20% sucrose. Coronal sections of 30 μm thickness were prepared using Leica CM1850 cryostat microtome and antigen retrieval was carried out by treating them at 95°C for 30 min in sodium citrate buffer pH 9.0. After blocking with 1% BSA, the sections were incubated with anti-CD31 mouse monoclonal antibody (1:200 dilution) conjugated with Alexa Fluor 594 (SantaCruz, sc-376764) and anti-*Pb*GAPDH rabbit polyclonal serum (1:100 dilution) or anti-mouse CD3 rat monoclonal antibody (Invitrogen, 14-0032-82; 1;100 dilution) for 16 h at 4°C. The sections were then treated with FITC-conjugated donkey anti-rabbit IgG (SantaCruz, sc-2090) or FITC-conjugated goat anti-rat IgG (SantaCruz, sc-2011; 1:200 dilution), followed by 4′,6-diamidino-2-phenylindole (DAPI) staining. Anti-mouse β-APP rabbit polyclonal antibody (Thermo Fisher Scientific, 51-2700) was used in 1:200 dilution. All the images were captured using Olympus IX83 microscope with DP73 high-performance camera.

### Heme, Hz, hemoglobin and hemopexin estimations

Free heme levels in the plasma samples of WT- and KO-infected mice were quantified using Hemin colorimetric assay kit (BioVision, K672) as per the manufacturer’s protocol. The assay is specific for free heme and it utilizes peroxidase activity of hemin to facilitate the conversion of a colorless probe to a strongly colored compound with absorbance at 570 nm. The quantification of free heme in the parasite lysates was carried out by resuspending the parasite pellets in 5 volumes of hypotonic lysis buffer containing 5 mM Tris pH 7.5 with protease inhibitors and incubating them in ice for 30 min. The lysates were then centrifuged at 20,000 *g* for 20 min, 4°C, and the supernatants obtained were used for free heme estimation as mentioned above for the plasma samples. The protein content of the hypotonic lysates was measured by Micro BCA protein assay kit (Thermo Scientific, 23235) and the free heme levels were expressed per mg of protein. The Hz content of the WT and KO parasite pellets was estimated as described earlier^80^. In brief, the parasite pellet was resuspended in 1 ml of 100 mM sodium acetate buffer, pH 5.0 and left at 37°C for overnight, followed by centrifugation at 10,000 *g* for 5 min. The resultant pellet was resuspended in 100 mM Tris buffer pH 8.0 containing 2.5% SDS and incubated at 37°C for 30 min, followed by centrifugation at 10,000 *g* for 5 min. The pellet obtained was washed once with 100 mM sodium bicarbonate pH 9.2 and then with distilled water. The final Hz pellet was dissolved in 100 mM NaOH containing 2.5% SDS, and the absorbance was measured at 405 nm. The supernatants of sodium acetate and Tris SDS steps were collected to estimate the protein content and heme content of the Hz was expressed per mg of total protein. To estimate the Hz content in the spleen and liver of the WT- and KO-infected mice, 50 mg tissue of the respective organs was homogenized in 50 mM Tris pH 8.0 containing 50 mM NaCl, 5 mM CaCl_2_ and 1% Triton X-100, and incubated for 12 h at 37°C in the presence of proteinase K. The lysates were sonicated and centrifuged at 15,000 *g* for 30 min. The pellets obtained were resuspended with 100 mM sodium bicarbonate containing 2% SDS and sonicated, followed by centrifugation at 15,000 *g* for 15 min. After repeating this step thrice, the pellets were solubilized in 100 mM NaOH containing 2% SDS and 3 mM EDTA, and the absorbance was measured at 405 nm^81^. In parallel, the parasite load in the organs was examined by quantitative PCR (qPCR) analysis carried out with *Pb18SrRNA* primers for the total RNA isolated from 30 mg tissue of the organs using RNeasy Plus Mini Kit (Qiagen, 74134). After normalizing with respect to the parasite load, the total heme content of the Hz isolated was expressed per mg weight of the organ. Hemoglobin and hemopexin levels in the plasma samples of WT- and KO-infected mice were measured by Mouse Hemoglobin (Abcam, ab157715) and Mouse Hemopexin (Abcam, ab157716) ELISA kits as per the manufacturer’s protocols.

### *In vivo* bioluminescence imaging

*In vivo* bioluminescence imaging for mice infected with WT and KO parasite lines expressing luciferase was carried out as described earlier^82^. Luc-expressing ALASKO and FCKO parasites were generated by transfecting WT parasites with GOMO-GFP-Luc plasmid containing GFP-Luc-expressing cassette with m-cherry flanked on either side by 5’- and 3’-UTR regions of the respective genes. The primers utilized were similar to the earlier ones^44^ except that the restriction sites used for forward and reverse primers were *SacII* and *NotI* for 5’ UTRs and *XhoI* and *KpnI* for 3’ UTRs, respectively. For Luc-expressing WT parasites, the following sets of forward and reverse primers were used to amplify the 5’- and 3’-UTRs of small subunit ribosomal RNA (*ssurRNA*): 5’UTR (F) – GCCACCGCGGGAAATACGACCAATATGTAATTATTGGATAATAATTG; 5’UTR (R) – GCCCGCGGCCGCCTACTGGCAAGATCAACCAGGTTACTATATATA; 3’UTR (F) – GCCACTCGAGGAGGCTTATCCTTCCTGATAAAGTG; 3’UTR (R) – GCCCGGTACCCAAAATACTAACCCACTATGTGCAATGTGC. For whole body imaging, 7-8 weeks old C57BL/6 mice were injected with Luc-expressing 10^5^ WT or 10^7^ KO parasites. On day 8 post-infection, mice were injected intraperitoneally with D-luciferin substrate (100 mg/kg animal weight in 200 μl of PBS), VivoGlo (Promega, P1041), and imaged after 5 min using *in vivo* Imaging System IVIS Lumina *XR* with medium binning, 10 sec exposure and 12.5 FOV, under XGI-8 gas anesthesia system. For *ex vivo* imaging, transcardial perfusion was carried out for infected mice with cold PBS after injecting D-luciferin, and the organs were dissected out and imaged.

### Analyses of cytokines, chemokines, chemokine receptors and other key mediators of cerebral pathogenesis

Bio-Plex assays for cytokines and chemokines were carried out in Bio-Plex 200 system using Bio-Plex Pro Mouse Cytokine Grp I Panel 23-Plex assay kit (Bio-Rad, M60009RDPD) following manufacturer’s protocol. The plasma samples utilized for the assays were prepared from the infected mouse blood collected on day7/8 post-infection. For transcript levels, total RNA was isolated from the brain samples of WT and KO-infected mice that were collected after a thorough perfusion with cold PBS. qPCR analyses were performed using QuantiFast SYBR Green RT-PCR Kit (Qiagen, 204154) on StepOne Real-Time PCR System (Applied Biosystems). The primers used were listed in Supplementary table 4. Expression levels were normalized with GAPDH and fold changes for the transcripts of KO-infected mice with respect to WT were calculated using 2^(-ΔΔ*Ct*)^ method.

### Flow cytometry

Flow cytometry analyses of T cells in the brain samples of WT- and KO-infected mice were carried out as described^83^. In brief, mice were anesthetized and transcardially perfused with PBS, and the brain samples were dissected out and harvested in RPMI-1640 medium containing 10% FBS. For preparing single cell suspensions, the samples were minced and digested in RPMI-1640 medium containing 0.05% Collagenase D and 2U/ml DNase I for 30 minutes at room temperature, and passed through 70 μm nylon cell strainer, followed by 5 minutes of incubation on ice. Brain homogenates were then overlaid on 30% Percoll cushion and centrifuged at 400 *g* for 20 minutes at room temperature. The leukocyte pellets obtained were resuspended in 1ml of RBC lysis buffer (155 mM NH_4_Cl, 10 mM NaHCO_3_ and 0.1 mM EDTA; pH 7.3) and incubated on ice for 5 minutes to remove any residual RBCs. The pellets were then washed with RPMI-1640, counted and stained for various markers. For intracellular markers like TNFα, IFNγ, perforin and granzyme B, staining was carried out after fixing the cells with 2% paraformaldehyde. Flow cytometry was performed with BD LSRFortessa and the data was analysed using FlowJo™ v10.6.1. The following fluorescent dye-conjugated antibodies were used for staining: anti-mouse CD3-FITC (clone 17A2; eBioscience), anti-mouse CD4-PE (clone RM4-5; eBioscience), anti-mouse CD8-PerCP-Cyanine5.5 (clone 53-6.7; eBioscience), anti-mouse CD69-Brilliant Violet 421 (clone H1.2F3; BioLegend), anti-mouse CXCR3-PE (clone CXCR3-173; eBioscience), anti-mouse Perforin-PE (clone S16009A; BioLegend), anti-mouse Granzyme B-APC (clone NGZB; eBioscience), anti-mouse IFNγ-eFluor 450 (clone XMG1.2; eBioscience), anti-mouse TNFα-eFluor 450 (clone MP6-XT22; eBioscience).

### FV isolation and analyses

In *P. berghei,* the Hz containing vacuoles tend to remain as small discrete vacuoles until the late schizont stage and they coalesce only at the time of schizont rupture. Because of this, purifying the FVs for *P. berghei* trophozoites is difficult using the standard percoll protocol that is followed for *P. falciparum,* wherein the small vacuoles coalesce in the late ring stages itself. Hence, we resorted ourselves to the isolation of FVs that are released during the schizont rupture as described with slight modifications^84^. In brief, the infected blood containing WT or FCKO parasites was collected around 22:00 h, centrifuged at 1,000 *g* for 5 min to remove plasma and buffy coat, and washed twice with RPMI-1640 medium containing 10% FBS. The washed cells were resuspended in 10 volumes of RPMI-1640 medium containing 10% FBS and then incubated at 37°C for overnight in a CO2 incubator. The maturation of schizonts was monitored by Giemsa smears and the FVs released in the culture supernatant during schizont rupture were collected in the next day around 09:00 h. In brief, the cultures were centrifuged at 200 *g* for 5 min to remove the RBCs. After repeating this step twice, the supernatant devoid of RBCs was centrifuged at 400 *g* for 10 min to collect the FVs that are free of merozoites. The FV pellet was then washed twice with PBS by centrifuging at 3000 *g* for 3 min and stored at -20°C. The purity of FVs was tested under microscope with 100x objective. Similar approach was followed for the isolation of FVs from griseofulvin-treated and -untreated parasites.

### Microscopy analyses of LysoTracker Deep Red uptake, BODIPY 493/503 and Nile Red staining, Hz content and Hz dynamics

To examine the LysoTracker Deep Red uptake in the live parasites, 10-20 μl of infected blood was collected from tail vein of the infected mice when the peripheral blood parasitemia was around 5-10% and resuspended in heparinised RPMI-1640 medium containing 10% FBS. After washing twice, the cells were incubated with 100 nM LysoTracker Deep Red (Thermo Fisher Scientific, L12492) for 30 min at 37°C in RPMI-1640 medium containing 10% FBS. The cells were then washed thrice with RPMI-1640 medium without phenol red, resuspended in the same medium and examined immediately under 100x objective using Olympus IX83 microscope with DP73 high-performance camera at 1600x1200 resolution using TRITC filter. BODIPY 493/503 (Thermo Fisher Scientific, D3922) and Nile Red (Sigma-Aldrich, 19123) staining for parasitized RBCs were carried out as described earlier^85^ In brief, 10-20 μl of infected mouse blood was collected from the tail vein in Hank’s balanced salt solution (HBSS) and centrifuged at 2000 *g* for 3 min. The cell pellet obtained was resuspended in 200 μl of HBSS containing BODIPY 493/503 (10 μg/ml) or Nile Red (2 μg/ml) and incubated at 37°C for 30 min. The cells were then washed twice and resuspended in HBSS. The images were acquired under 100x objective using Olympus IX83 microscope with DP73 high-performance camera at 1600x1200 resolution using FITC/TRITC filters. The images for WT and KO were acquired under identical exposure conditions and the fluorescent signal intensities were quantified using ImageJ software. To examine the Hz content, bright field and DIC images were taken under 100x objective. Live imaging of Hz dynamics was carried out under 100x objective by acquiring 15 frames per second at 1600x1200 resolution. The video files were processed using VSDC video editor and VideoPad by NCH softwares.

### Labelling studies with ^14^C-ALA, ^32^P-orthophosphoric acid and ^14^C-SA

[4-^14^C]-ALA (ARC 1550), ^32^P-orthophosphoric acid (ARC 0103) and [1-^14^C]-SA (ARC 025) were procured from American Radiolabeled Chemicals, Inc. The infected blood samples were collected, centrifuged at 1,000 *g* for 5 min to remove plasma and buffy coat, and washed twice with RPMI-1640 medium containing 10% FBS. The washed cells were resuspended in 10 volumes of RPMI-1640 medium containing 10% FBS and then incubated at 37°C in a CO_2_ incubator with the respective radioactive compounds. For ^14^C-ALA labelling, blood samples were collected from griseofulvin treated and control WT-infected mice around 16:00 h and the labelling was carried out for 9 h at a radioactivity of 1 μCi/ml. The infected RBCs were then centrifuged, washed with PBS and the parasites were isolated by saponin treatment. After washing the parasite pellet with PBS for four times, free heme present in the parasites was extracted using ethylacetate:glacial acetate (4:1) followed by 1.5 N HCl and water washes to remove porphyrins and ALA as described earlier^44^. The upper phase was separated, dried under nitrogen stream, dissolved in methanol and analysed by TLC on silica gel using the mobile phase 2,6-lutidine and water (5∶3 v/v) in ammonia atmosphere. The intensity of radiolabelling was scanned using Amersham Typhoon 5 Biomolecular Imager by exposing the TLC sheets to phosphorimager screen and the radioactive counts were measured using PerkinElmer MicroBeta^2^ 2450 Microplate Counter. For ^32^P-orthophosphoric acid and ^14^C-SA labelling, the blood sample collection and incubation were carried out as mentioned for the FV isolation. In addition to the isolation of the secreted FVs, the infected RBC pellets were subjected to saponin treatment for the isolation of the radiolabelled total parasites. The labelling was carried out for ^32^P-orthophosphoric acid and ^14^C-SA at radioactivities of 50 μCi/ml and 2 μCi/ml, respectively. All these experiments were typically carried out in a total volume of 4 ml, carefully matching in terms of parasitized RBCs and haematocrit between WT and FCKO.

### Lipid analyses

Lipid extraction for phospholipid analysis was carried out as described^86^. In brief, the FV and parasite pellets were extracted with chloroform:methanol (2:1) and washed twice with water. The lower organic phase was collected and analysed by TLC on silica gel using the mobile phase chloroform:ethanol:water:tri-ethylamine (30:35:7:35 v/v). 5 μl of the sample was used to take radioactive counts. For fatty acid analysis, the lipids extracted as mentioned above were dried, dissolved in 50 μl of toluene and subjected to mild methanolysis/methylation by the addition of 375 μl of methanol and 75 μl of 8% HCl in methanol. After incubating at 45°C for 16 h, FAMEs were extracted by the addition of 250 μl of hexane and 250 μl of water^87^. The hexane layer containing FAMEs was analysed by TLC on silica gel impregnated with 0.5% methanolic silver nitrate using the mobile phase light petroleum ether:acetone:formic acid (97:2:1 v/v) to separate FAMEs derived from unsaturated fatty acids^88^. In this separation, SAME lacking double bond migrates faster as an uppermost top band followed by OAME and other unsaturated FAMEs that are separated based on their degree of unsaturation and the length of fatty acyl chain. The respective standards were used for all the lipids. 5 μl of the hexane layer was used to take radioactive counts. For GC-MS analysis, 1 µl of hexane containing the FAMEs was injected in split mode (50:1). The GC-MS analysis was performed using Agilent 7890B GC coupled with 240 Ion Trap MS. The column used for analysis was Agilent VF-5MS (30 m length x 0.25 mm internal diameter (ID) x 0.25 µm film thickness). The oven temperature was programmed from 140°C (5 min hold), increased at rate of 4°C/min to 240°C. Helium was used as carrier gas with 1 ml/min flow rate. The mass spectrometer was operated in full scan mode from 40 to 500 m/z and NIST library search was performed to identify the compounds.

### Proteomics analyses

To examine the protein content of the FVs from WT and FCKO parasites, proteins were extracted from three different preparations of WT and FCKO FVs using 25 mM ammonium bicarbonate containing 6M urea, followed by treatment with DTT and iodoacetamide. The urea concentration was reduced to 0.8 M by performing dilution with 25 mM ammonium bicarbonate. In-solution trypsin (SCIEX) digestion was carried out for 200 μg total protein, followed by desalting. LC-MS/MS analyses were performed using SCIEX TripleTOF 5600+ ESI-Mass Spectrometer. The acquired data were analysed using Paragon algorithm against *Plasmodium berghei* Fasta files from UniProt for 95% confidence.

### Griseofulvin treatment in mice

Griseofulvin (Sigma-Aldrich) was prepared by dissolving 1 or 2 mg in 40 μl DMSO and then making up the volume to 200 μl with 10% solutol HS 15 (Sigma-Aldrich) in saline. The mixture was vortexed thoroughly for 10 min to form an emulsion and injected intraperitoneally into the mice. All single dose injections were carried out at 06:00 h and double dose injections were carried out at 06:00 h and 18:00 h for the respective days. Control mice were injected with the solvent.

### Western blot analyses and other procedures

Western analyses were carried out using standard procedures. In brief, brain samples were homogenized in 50 mM Tris-Cl buffer, pH 7.5 containing 5 mM EDTA, 50 mM NaCl, 5 mM DTT, 0.1% Np-40, 50 mM NaF, 1 mM PMSF, 1 mM Na_3_VO_4_, and 1x Halt Protease Inhibitor Cocktail (Thermo Fisher Scientific), followed by centrifugation at 18,000 *g* for 20 min at 4°C. The supernatants were collected and quantified for total protein. The following antibodies were used: anti-mouse NF-κB p65 (Invitrogen, 14-6731-81); anti-mouse Phospho-NF-κB p65 (Ser536) (Invitrogen, MA5-15160); anti-mouse NLRP3 (Invitrogen, PA5-20838); anti-mouse Phospho-NLRP3 (Ser295) (Invitrogen, PA5-105071); anti-mouse Caspase 1 (Invitrogen, 14-9832-82); anti-mouse Cleaved Caspase-1 (Asp296) (Cell Signaling Technology, 89332); anti-mouse IL-1β (Invitrogen, # 701304); anti-mouse Cleaved IL-1β (Cell Signaling Technology, 52718) and anti-mouse β-Actin (Cell Signaling Technology, 3700). Blots were developed using Pierce ECL Western Blotting Substrate (Thermo Fisher Scientific).

### Statistical analyses

All the graphs were plotted using GraphPad Prism 7 software and the statistical analyses were carried out using unpaired t-test and log-rank (Mantel-Cox) test. n.s - not significant, \**P*<0.05, \*\**P*<0.01, \*\*\**P*<0.001.

## SUPPLEMENTARY INFORMATION

**Supplementary Figure 1:**
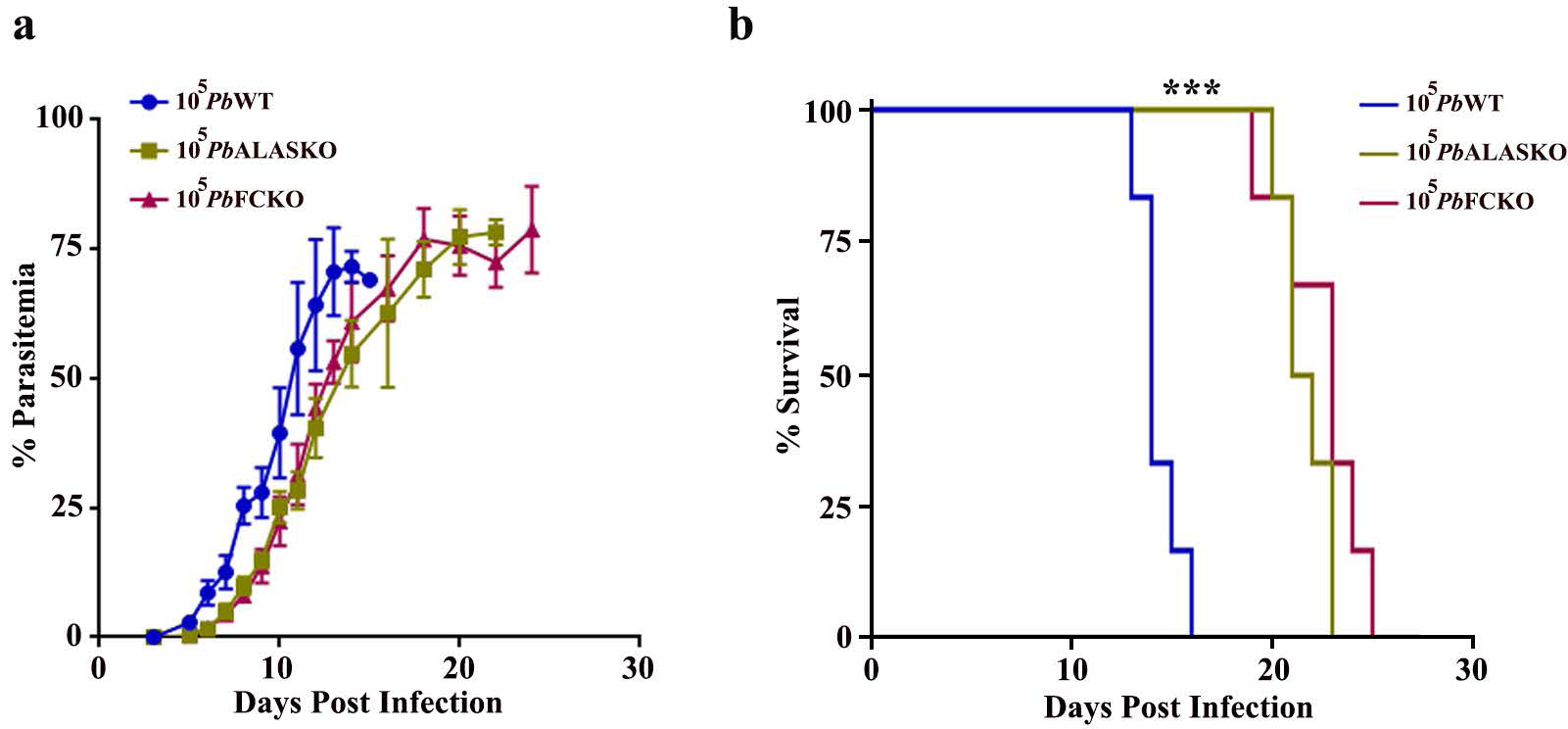
Growth characterization of heme pathway KO parasites in Balb/c mice. **a**, Growth analysis in Balb/c mice (n=6). 10^5^ parasites were used to initiate parasite infections. **b**, Mortality curves (n=6) (\*\*\**P*<0.001, log-rank (Mantel-Cox) test).

**Supplementary Figure 2:**
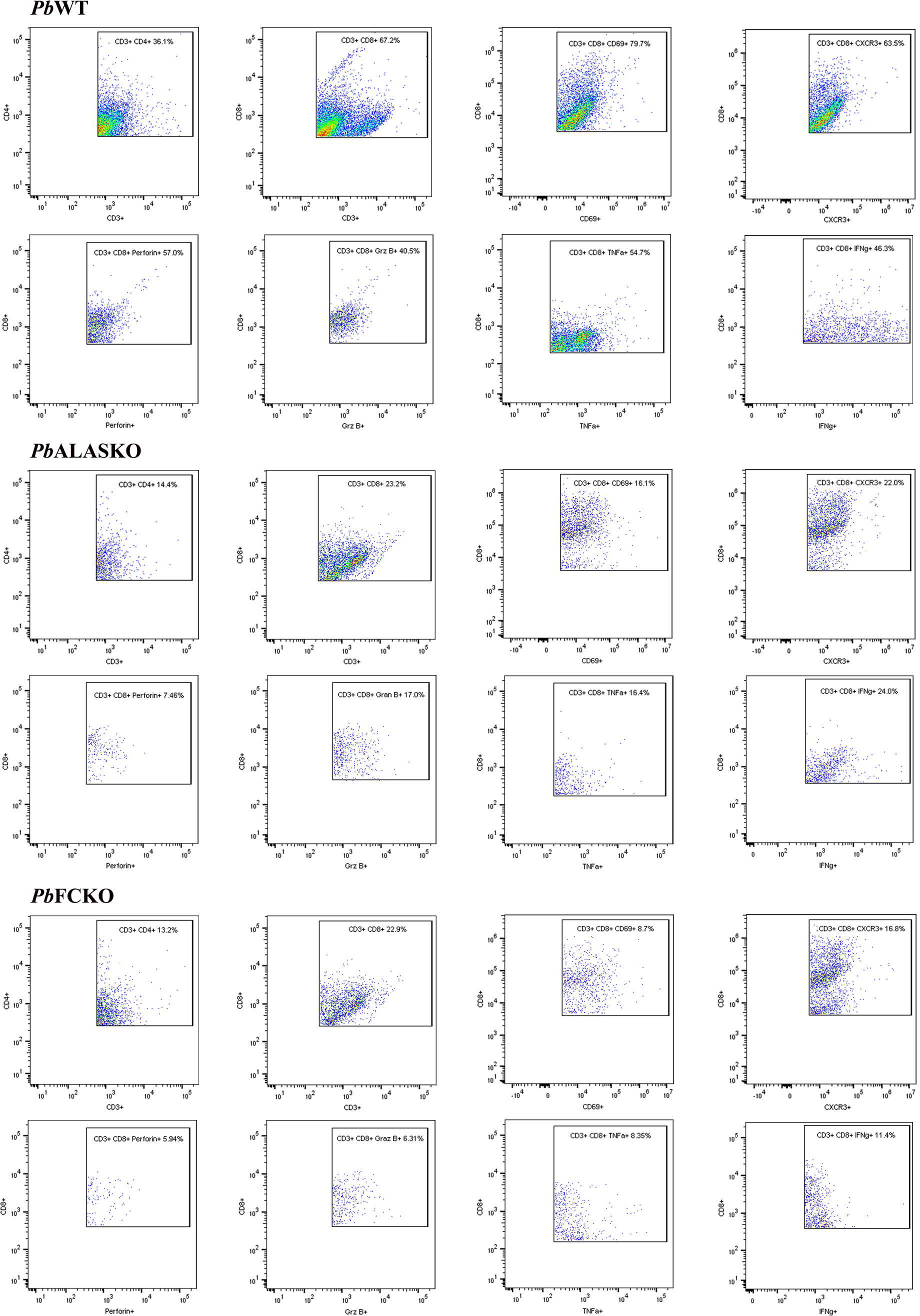
Dot plots from flow cytometry analyses representing the T cells in the brain samples of WT and KO-infected mice. The data represent CD3^+^ CD4^+^ and CD3^+^ CD8^+^ double positive cells, and CD3^+^ CD8^+^ CD69^+^, CD3^+^ CD8^+^ CXCR3^+^, CD3^+^ CD8^+^ perforin^+^, CD3^+^ CD8^+^ granzyme B^+^, CD3^+^ CD8^+^ TNFα^+^ and CD3^+^ CD8^+^ IFNγ^+^ triple positive cells, obtained for WT-, ALASKO- and FCKO-infected mice.

**Supplementary Figure 3:**
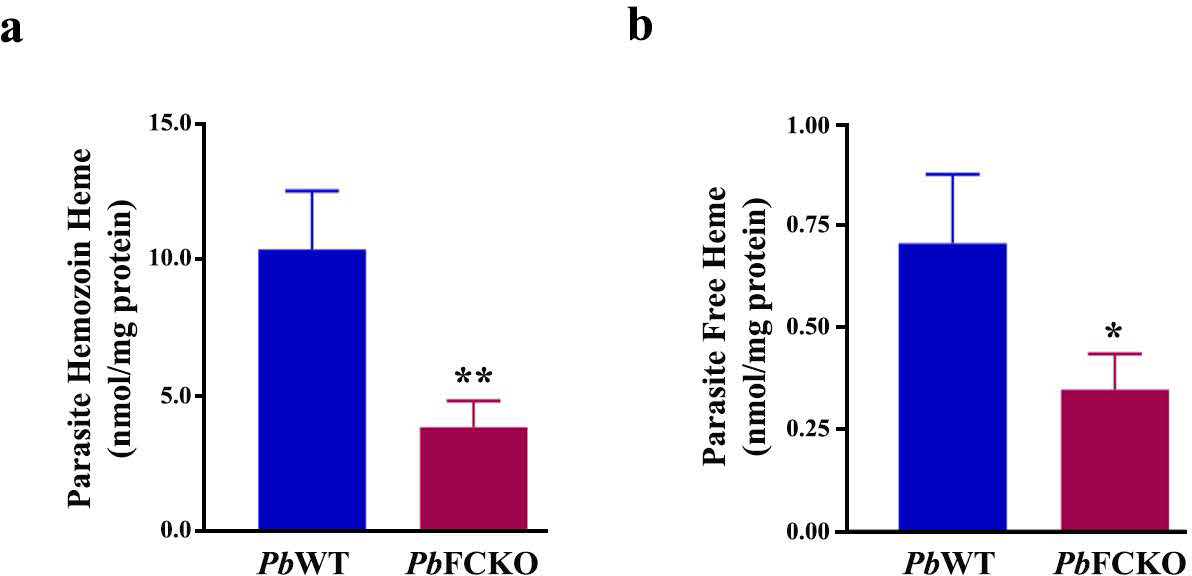
Hz and free heme levels in WT and FCKO parasites isolated from Balb/c mice. **a,b**, Hz and free heme levels in WT and FCKO parasites (n=3), respectively (mean ± SD \**P*<0.05, \*\**P*<0.01, unpaired t-test).

**Supplementary Figure 4:**
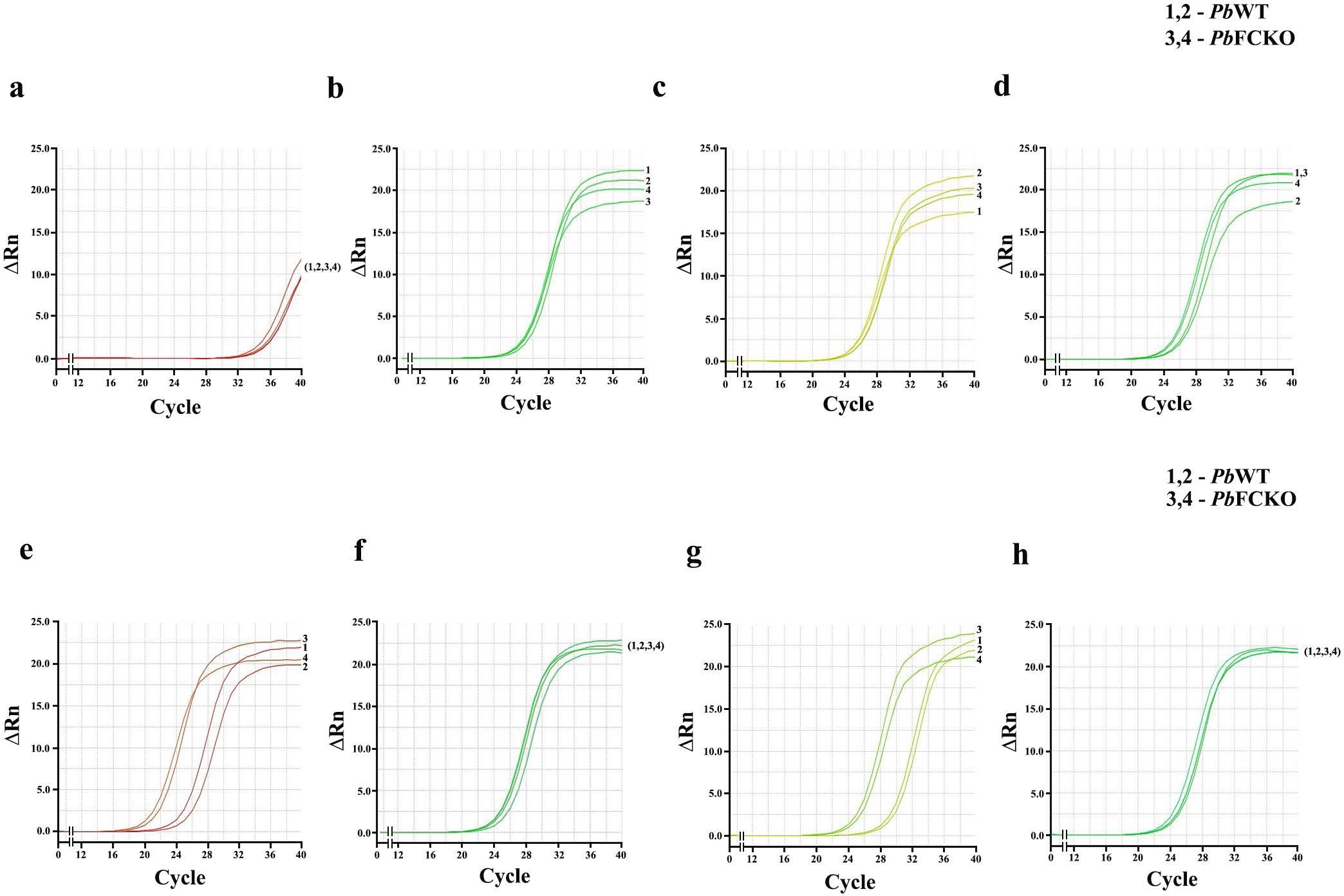
Assessment of parasite load in α,β-arteether and chloroquine treated WT- and KO-infected mice. **a**, qPCR analyses of parasite load using *Pb*GAPDH primers for the infected mice treated with 1 mg/mouse dose of α,β-arteether with total RNA isolated from the whole blood. The amplification curves represent primer dimers indicating the absence of detectable parasites. **b**, qPCR analyses for mouse GAPDH control. **c**, qPCR analyses of parasite load using *Pb*GAPDH primers for the infected mice treated with 0.4 mg/mouse dose of α,β-arteether. **d**, qPCR analyses for mouse GAPDH control. The Δ*C_t_* value obtained for FCKO with respect to WT for 0.4 mg dosage was -1.43 ± 0.16 (mean ± SD). **e**, qPCR analyses of parasite load using *Pb*GAPDH primers for the infected mice treated with 25 mg/kg dose of chloroquine. **f**, qPCR analyses for mouse GAPDH control. **g**, qPCR analyses of parasite load using *Pb*GAPDH primers for the infected mice treated with 10 mg/kg dose of chloroquine. **h**, qPCR analyses for mouse GAPDH control. The Δ*C_t_* values obtained for FCKO with respect to WT were 4.02 ± 0.91 and 4.19 ± 0.43 for 25 mg/kg and 10 mg/kg doses, respectively (mean ± SD). The *C_t_* values obtained for mouse GAPDH were comparable between WT- and FCKO-infected mice. 1,2 - WT-infected mice; 3,4 - FCKO-infected mice.

**Supplementary Figure 5:**
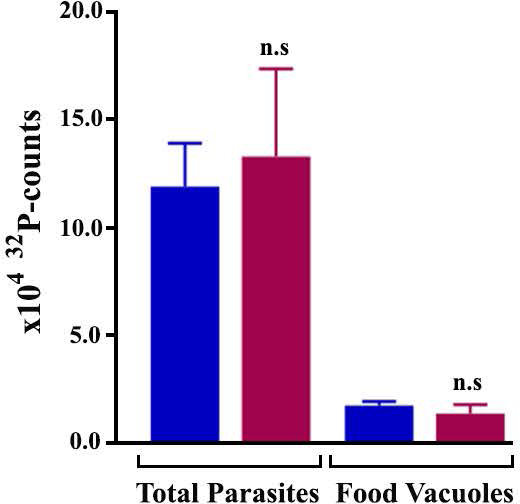
Radiolabelling of phospholipids in WT and FCKO parasites. Radioactive counts for ^32^P-orthophosphoric acid radiolabelled phospholipids of WT and FCKO total parasites and FVs. The data represent three different experiments (mean ± SD; n.s - not significant, unpaired t-test).

**Supplementary Figure 6:**
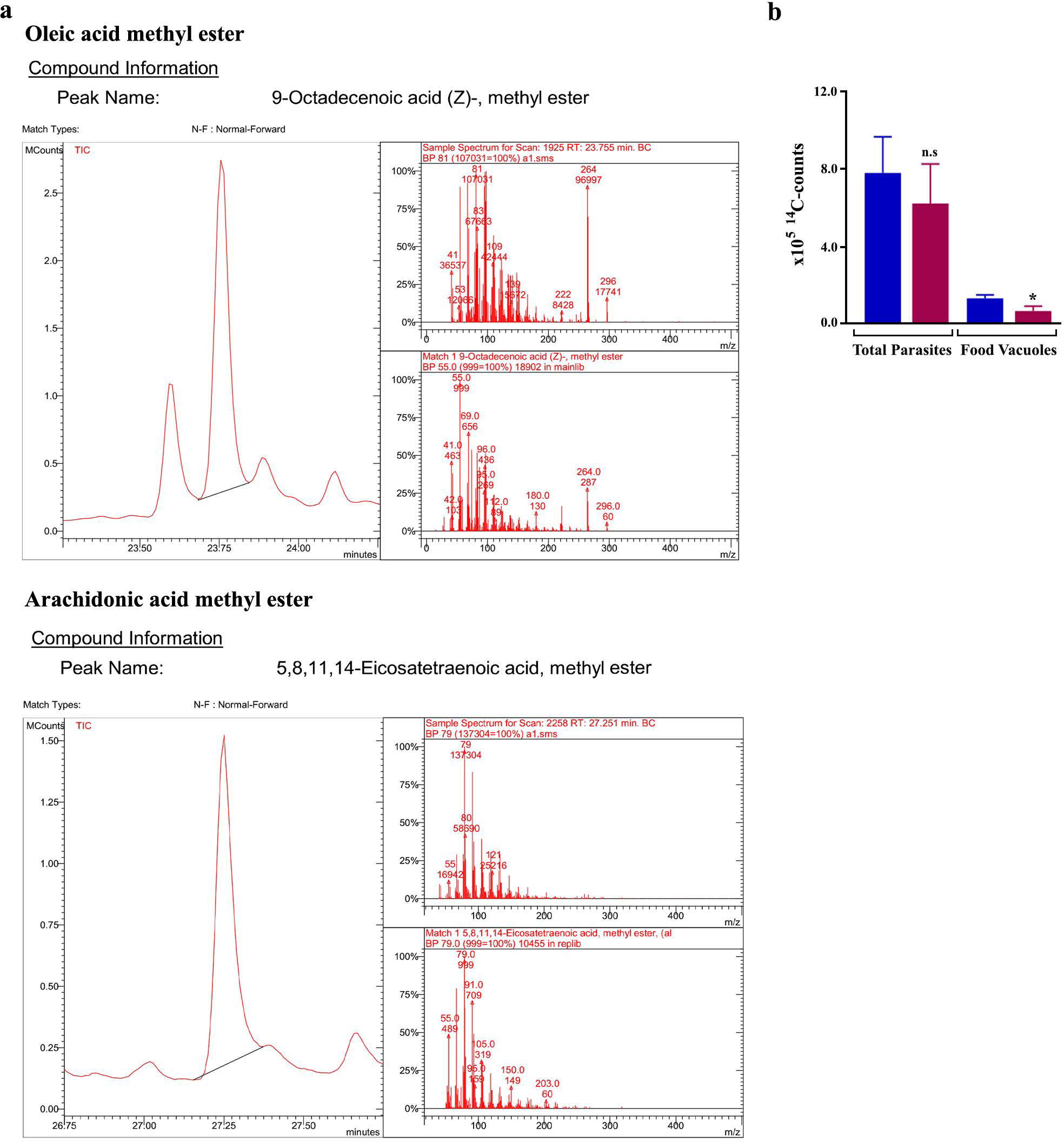
FAMEs prepared from *Pb* FVs. **a**, GC-MS analysis of FAMEs prepared from *P. berghei* FVs. The chromatogram peaks for the methyl esters of oleic acid and arachidonic acid and their mass spectra are shown. The entire set of compounds that could be identified and their respective area of peaks are given in Table S2. **b**, Radioactive counts for ^14^C-SA radiolabelled FAMEs of WT and FCKO total parasites and FVs. (mean ± SD; n.s - not significant, *P<0.05, unpaired t-test)

**Supplementary Table 1: List of compounds identified in GC-MS analysis of the FAMEs prepared from *Pb* FVs.** Methyl esters of oleic acid and arachidonic acid are highlighted in green. The compounds are listed in the order of their retention time.

**Supplementary Table 2: *Pb*WT FV proteome and gene ontology clustering.** Proteins that are identified only in WT FVs, but not in FCKO FVs are highlighted in green. The entire peptide summary along with the identified sequences is also provided.

**Supplementary Table 3: *Pb*FCKO FV proteome and gene ontology clustering.** Proteins that are identified only in FCKO FVs, but not in WT FVs are highlighted in blue. The entire peptide summary along with the identified sequences is also provided.

**Supplementary Table 4:** Details of the primers used for qPCR analyses of the host transcripts in the brain samples of WT- and KO-infected mice.

**Supplementary Movie 1: Live imaging of hemozoin dynamics in *Pb*WT parasite.**

**Supplementary Movie 2: Live imaging of hemozoin dynamics in *Pb*ALASKO parasite.**

**Supplementary Movie 3: Live imaging of hemozoin dynamics in *Pb*FCKO parasite.**

## Notes

### Competing Interest Statement

The authors have declared no competing interest.

